# Phylogenetic, population genetic, and morphological analyses reveal evidence for one species of Eastern Indigo Snake (*Drymarchon couperi*)

**DOI:** 10.1101/318766

**Authors:** Brian Folt, Javan Bauder, Stephen Spear, Dirk Stevenson, Michelle Hoffman, Jamie R. Oaks, Christopher Jenkins, David A. Steen, Craig Guyer

**Affiliations:** Department of Biological Sciences and Auburn University Museum of Natural History, 331 Funchess Hall, Auburn University, Alabama 36849, U.S.A.; The Orianne Society, 11 Fruitstand Lane, Tiger, Georgia 30576, U.S.A.; Department of Environmental Conservation, University of Massachusetts, Amherst, Massachusetts, U.S.A.; The Wilds, Cumberland, Ohio, U.S.A.; The Orianne Center for Indigo Conservation, Central Florida Zoo and Botanical Gardens, 3755 NW Hwy 17-92, Sanford, Florida, 32771, U.S.A.

## Abstract

Accurate species delimitation and description are necessary to guide effective conservation management of imperiled species. The Eastern Indigo Snake (*Drymarchon couperi*) is a large species in North America that is federally-protected as Threatened under the Endangered Species Act. Recently, two associated studies hypothesized that *Drymarchon couperi* is two species. Here, we use diverse approaches to test the two-species hypothesis for *D. couperi*. Our analyses reveal that (1) phylogenetic reconstruction in previous studies was based entirely on variance of mitochondrial DNA sequence data, (2) microsatellite data demonstrate significant population admixture and nuclear gene flow between mitochondrial lineages, and (3) morphological analyses recover a single diagnosable species. Our results are inconsistent with the two-species hypothesis, thus we reject it and formally place *Drymarchon kolpobasileus* into synonymy with *D. couperi*. We suggest inconsistent patterns between mitochondrial and nuclear DNA may be driven by high dispersal of males relative to females. We caution against species delimitation exercises when one or few loci are used without evaluation of contemporary gene flow, particularly species with strong sex-biased dispersal (e.g., squamates) and/or when results have implications for ongoing conservation efforts.

## Introduction

Accurate species delimitation and description are critical not only for understanding global patterns of biodiversity, but also to guide effective conservation strategies [1–4]. For example, species are often delimited into multiple species on the basis of systematic studies utilizing molecular genetic data, thereby requiring adjustment of existing conservation management plans (e.g., [5]). When species delimitation methods fail to correctly diagnose individuals (*sensu* [6]), such errors can have significant consequences for conservation and management of imperiled species by reducing or diverting finite conservation resources [2]. Therefore, taxonomic division into multiple species should be performed carefully and only when robust evidence supports a decision to revise. Indeed, authors have cautioned that studies of species delimitation should be conservative, because “it is better to fail to delimit species than it is to falsely delimit entities that do not represent actual evolutionary lineages” [7].

The Eastern Indigo Snake (*Drymarchon couperi*) is a large colubrid native to the Coastal Plain of the southeastern United States. However, *D. couperi* populations have declined in abundance precipitously over the last century, largely due to habitat loss, habitat fragmentation, and historical over-collecting for the pet trade [8,9]. As a result of these declines, *D. couperi* is listed as Threatened under the U.S. Endangered Species Act [9,10]. Potentially viable populations of *D. couperi* remain in large contiguous habitats in southeastern Georgia [11–13], and throughout peninsular Florida [13,14], but the species was likely extirpated from Mississippi, Alabama, and the Florida panhandle [13].

Current conservation management plans for *D. couperi* were developed under the hypothesis that *D. couperi* represents a single species. However, this hypothesis was recently challenged by Krysko et al. [15], who used DNA sequence analyses to describe two genetic lineages of *D. couperi* – an Atlantic lineage, including populations in southeastern Georgia and eastern peninsular Florida, and a Gulf lineage of populations in western and southern peninsular Florida and the Florida panhandle. This phylogeographic study was followed by a second paper [16] that analyzed morphological variation between the Atlantic and Gulf lineages and provided an official description of the Gulf lineage as a purported novel species, the Gulf Coast Indigo Snake (*Drymarchon kolpobasileus*).

Given the conservation status of *D. couperi* (*sensu lato*), these results have potentially important consequences for the conservation of Eastern Indigo Snakes. First, division of *D. couperi* (*sensu lato*) into two smaller-ranged species results in two species with substantially smaller population sizes that are, therefore, at greater risk of extinction (*sensu* [2]; e.g., [17]). Second, conservation and recovery of two rare species requires more time and funds than one, and both resources are in short supply. Finally, as noted by Krysko et al. [15], active conservation management plans for *D. couperi* (*sensu lato*) include population repatriation projects in Alabama and the Florida panhandle, where populations attributed to the Gulf lineage presumably were extirpated. Repatriation projects should be informed by phylogeographic and genetic developments [18]. The description of *D. kolpobasileus*, therefore, causes increased logistical complexity for Eastern Indigo Snake captive breeding and repatriation projects.

We represent additional experts on Eastern Indigo Snake taxonomy, ecology and conservation, many of whom participated in an inter-agency workshop on Eastern Indigo Snake taxonomy referenced in Krysko et al. [15,16]. During that event, consequences of the discovery that Eastern Indigo Snakes comprise two genetic lineages were debated, and this debate was used to inform conservation plans for the species. However, skepticism was voiced that the two lineages represent distinct evolutionary species, based largely on description of microsatellite data documenting widespread admixture of the lineages. Here, we formally address that debate to evaluate the hypothesis that *D. couperi* comprises two species [15,16]. We adopted a unified species concept [19,20] to define and operationally diagnose species. Under this concept, species are necessarily defined as independently evolving metapopulation lineages and this feature must necessarily be demonstrated to delimit species [20]. However, we also evaluated additional lines of evidence, such as morphological diagnosability, to provide secondary assessments of lineage separation. Together, we sought to provide a conceptually robust and integrative test [21,22] of whether *D. couperi* is two distinct species.

In this paper, we first re-analyze gene sequence data presented by Krysko et al. [15] to infer whether their phylogeny was overly-influenced by data from the mitochondrial genome. Second, we analyze a novel microsatellite DNA dataset and test for evidence of population admixture and contemporary gene flow between the two genetic lineages identified by Krysko et al. [15] as an explicit test of whether there are two independently evolving metapopulation lineages of *D. couperi*. Specifically, if *D. couperi* is two species, we predicted little or no admixture between hypothetical species, particularly at the putative contact zone identified by Krysko et al. [15,16]. Third, we analyze new morphological data collected from 125 Eastern Indigo Snakes, including individuals from both genetic lineages of Krysko et al. [15,16]. Specifically, we evaluate the diagnostic features of head and scale shape presented by Krysko et al. [16] as tests of whether morphological phenotypes are secondary lines of evidence supporting previously delimited species. Last, we review features of the life history of *D. couperi* and suggest how they inform interpretations of genetic and morphological data.

## Materials and Methods

### Gene sequence analysis

To infer evolutionary history among populations of *D. couperi* (*sensu lato*), Krysko *et al.* [15] analyzed sequence data obtained from three genetic markers: the linked mitochondrial (mtDNA) genes cytochrome *b* (CytB) and nicotinamide adenine dinucleotide dehydrogenase subunit 4 (ND4) and the nuclear gene *neurotrophin*-3 (NT3). The authors estimated phylogenetic relationships among populations by analyzing a concatenated dataset including both mitochondrial and nuclear loci. These data were evaluated with maximum likelihood (ML) and Bayesian analyses; for the Bayesian analysis, the dataset was partitioned such that nucleotide substitution was modeled separately for each locus. Because both analyses generated similar phylogenetic hypotheses, the authors described results only from the Bayesian analysis.

Phylogenetic analyses of a single or few genetic loci frequently describe evolutionary patterns that do not reflect the organism’s true evolutionary history (i.e., the gene tree/species tree problem; [1,23–25]). In particular, use of and reliance on mtDNA for phylogenetic and taxonomic analyses has been criticized because mtDNA has a vastly different natural history than the primary mode of genetic inheritance, nuclear DNA (nDNA). Mitochondrial DNA has a lower effective population size, higher mutation rate, and frequently defies critical assumptions of neutral evolution by being under selection [4,26]. More importantly, mtDNA is maternally inherited and, therefore, may not describe an organism’s true patterns of inheritance expressed through the nuclear genome [26]. This is particularly problematic for species with relatively low dispersal rates that are more likely to show phylogeographic breaks that are not driven by decreased gene flow but by chance alone [27], or for species with intersexual differences in movement, site fidelity, or breeding behavior [28–30].

Given these and other limitations, a customary practice in phylogenetic studies is to use both mitochondrial and nuclear loci and to describe phylogenetic patterns inferred from these two components of the genome separately (e.g., [24,31–33]). This practice can help identify situations for which phylogenetic hypotheses generated from mtDNA (1) are incongruent with hypotheses from the nuclear genome and that (2) might be erroneously assumed to accurately depict the species tree. However, Krysko *et al.* [15] combined the mitochondrial and nuclear markers and used that concatenated dataset to infer both ML and Bayesian phylogenies from the combined datasets. We therefore argue that the DNA sequence analyses of Krysko et al. [15] are biased toward describing patterns from maternally inherited mtDNA and require re-evaluation.

To explore the extent to which nuclear sequence data support lineage divergence and speciation within Eastern Indigo Snakes, we accessed the Krysko *et al.* [15] sequence data from GenBank (S1 Table) and used ML methods to infer a nuclear gene tree from the NT3 dataset, following the methods used by Krysko *et al.* [15]. This dataset included 23 *D. couperi* (*sensu lato*) samples (N = 13 Atlantic clade, N = 10 Gulf clade) and four outgroup taxa (*Drymarchon melanurus erebennus*, *Drymarchon melanurus rubidus*, *Coluber constrictor*, *Masticophis flagellum*). We fit different models of nucleotide substitution and ranked them using BIC; this procedure suggested that the Kimura [34] model best fit the data. We then estimated a ML phylogeny for NT3 using the package ‘phangorn’ [35] in the statistical Program R [36]. This method allowed us to test whether a phylogenetic pattern inferred from nuclear DNA alone was consistent with or different from a concatenated analysis of mitochondrial and nuclear DNA.

### Microsatellite analysis

To test whether mitochondrial lineages of *D. couperi* represent independently evolving metapopulation lineages, we extracted and genotyped microsatellite DNA from 428 tissue samples (scale, shed skins, or muscle from road-killed individuals) throughout peninsular Florida and southern Georgia. Twenty-five samples were obtained from the collections of the Florida Museum of Natural History, including 20 samples used in Krysko *et al.* [15]. The samples from Krysko *et al.* [15] included individuals from central Florida that represented both mitochondrial clades where they occur in close proximity. The remaining Florida samples (N = 170) were collected during field studies of *D. couperi* [37,38] in and around Highlands County or opportunistically by authorized project partners. The samples from Georgia (N = 233) were collected by multiple project partners as part of a study of population fragmentation in the state (S. Spear et al., unpublished data). Our samples include similar representation of both mitochondrial lineages (55% Atlantic and 45% Gulf). We extracted DNA using the Qiagen DNeasy blood and tissue extraction kit (Qiagen, Inc., Valencia, CA). We ran 17 microsatellite loci [39] within three multiplexed panels using the Qiagen Multiplex PCR kit (S2 Table for details). Each reaction contained 1X Qiagen Multiplex PCR Master Mix, 0.2 μM multiplexed primer mix (each primer at equal concentrations), and 1 μl of DNA extract in a total volume of 7 μl. The PCR protocol was modified from Shamblin et al. [39] for multiplex PCR and consisted of an initial denaturation of 95°C for 15 min, 20 touchdown cycles of 94°C for 30 s, 60°C minus 0.5°C per cycle for 90 s and 72°C for 1 min, followed by 30 cycles of 94°C for 30 s, 50°C for 90 s and 72°C for 1 min, and a final elongation step of 60°C for 30 min. Multiplexed PCR products were run on a 3130xl Applied Biosystems Genetic Analyzer at the University of Idaho’s Laboratory for Ecological, Evolutionary, and Conservation Genetics. We scored fragment sizes using Genemapper 3.7 (Applied Biosystems).

We tested for the presence of null alleles that would lead to violations of Hardy-Weinberg equilibrium assumptions using the software FreeNA [40] and excluded any loci that had an estimated null allele frequency > 0.10. We estimated population structure and number of genetic clusters using the Bayesian clustering algorithm Structure 2.3.4 [41]. We used the admixture model with 100,000 iterations following 10,000 burn-in repetitions. We evaluated K = 1–10 with five replicates for each value of K. We used Structure Harvester [42] to implement the Delta K method of Evanno et al. [43] to estimate the number of clusters that best explain the microsatellite data. We used CLUMPP v.1.1.2. [44] to estimate the optimal cluster assignment in a single file based on the five replicates for the best supported values of K. Because the Evanno et al. [43] method analyzes changes in likelihood between values of K, it cannot estimate Delta K for K = 1; therefore, we assessed the probability of a K = 1 scenario with the raw likelihood values from Structure. Given biases of methods to estimate population structure from microsatellite data [45], we sought to follow recommendations from Janes et al. [45] by describing and comparing population structure predicted by both the Delta K and raw likelihood outputs from Structure, while also reporting bar plot outputs for different values of K supported by those methods. We used the program CLUMPAK [46] to visualize bar plot outputs from Structure. We note that during these exploratory hierarchical analyses we observed additional fine scale genetic structure among populations; these details were outside the scope of the current analysis, but we intend to examine these data more fully in a future paper. Finally, we tested for evidence of population differentiation in the 20 samples used by Krysko *et al.* [15] that represented spatial overlap of Atlantic and Gulf clades by estimating Jost’s D metric of genetic differentiation [47]. Jost’s D was developed to better represent actual levels of genetic differentiation when markers with high mutation rates (such as microsatellites) are used. We estimated Jost’s D using the ‘mmod’ package [48] in R.

We also conducted a spatial principle components analysis (sPCA) to identify spatial patterns of genetic structure while accounting for spatial autocorrelation among samples without relying on assumptions of Hardy-Weinberg equilibrium [49]. sPCA requires specifying a connection network to define connected samples. We evaluated three connection networks: (1) Delaunay triangulation, (2) Gabriel graph, and (3) a distance-based connection network, where samples ≤ 22.2 km (the maximum known dispersal distance by Eastern Indigo Snakes; [50]) were considered connected. We conducted significance tests for global and local structure using 9,999 permutations in the package ‘adegenet’ [51] in R.

Last, we analyzed the microsatellite data using linear mixed-effects models with maximum-likelihood population effects (MLPE; [52]) to better understand the role of isolation by distance in explaining genetic distance within and among clades. We estimated genetic distance at the individual level using 1 - proportion of shared alleles [53], where increased values indicated greater genetic dissimilarity between samples. To test for isolation by distance, we built three MLPE models examining how genetic distance varied by Euclidean geographic distance: a model using only Atlantic clade samples, a model using only Gulf clade samples, and a model using all sample from both clades. For each model, we report the parameter estimate for Euclidean distance, it’s standard error and *t* statistic, and the marginal *R^2^* (i.e., the proportion of variance explained by fixed-effect factors; [54,55]). To visualize how isolation by distance differed within and between clades, we graphically overlaid genetic distance against Euclidean distance for within-clade distances against genetic distances measured between clades to see how isolation by distance varied within clades relative to across all samples. If mitochondrial lineages represented different species, we predicted to observe lower genetic distances within clades than between clades (i.e., no or little overlap in genetic distances within and among clades). We estimated proportion of shared alleles using the package ‘adegenet’ [51] and implemented MLPE tests with individuals as random effects using the package ‘ResistanceGA’ [56] in R.

### Morphological analysis

Krysko et al. [16] performed principal components analyses (PCA) on five linear measurements of head morphology: head length, head height, length of a temporal scale, length of the 7^th^ infralabial, and width of the 7^th^ infralabial. Head length, head height, and 7^th^ infralabial length loaded positively and 7^th^ infralabial width loaded negatively on the first principal component, which explained 44.7% of variance. This axis separated specimens that were described to be long‐ and wide-headed with long and narrow 7^th^ infralabials (high PC1 scores; almost exclusively Atlantic lineage snakes) from specimens that were described to be short and narrow-headed with short and wide 7^th^ infralabials (low PC1 scores; almost exclusively Gulf lineage snakes). Temporal length loaded heavily and positively on the second principal component, which explained 25.4% of variance and separated specimens with relatively elongate temporals (high values on PC2; primarily Atlantic lineage) from specimens with relatively short temporals (low values on PC2; primarily Gulf lineage). The authors then used these axes to justify presentation of the shape of the 7^th^ infralabial, temporal, and head as being diagnostic of each species.

We found inconsistencies in the head scale and shape characters measured by Krysko et al. [16], which caused us to question their validity as diagnostic features. In particular, the specific temporal scale identified as being measured differed between Figures 3 and 5 of Krysko et al. [16]. This resulted because the number and position of temporal scales within species of *Drymarchon* is variable (Fig 1). Similarly, Figure 3 of Krysko et al. [16] appears to identify the 6^th^ infralabial rather than the 7^th^ as the scale measured for multivariate analysis. Thus, it is unclear which scale was measured for this important character and whether the same scale was measured among all individuals included in analysis.

**Fig 1.**
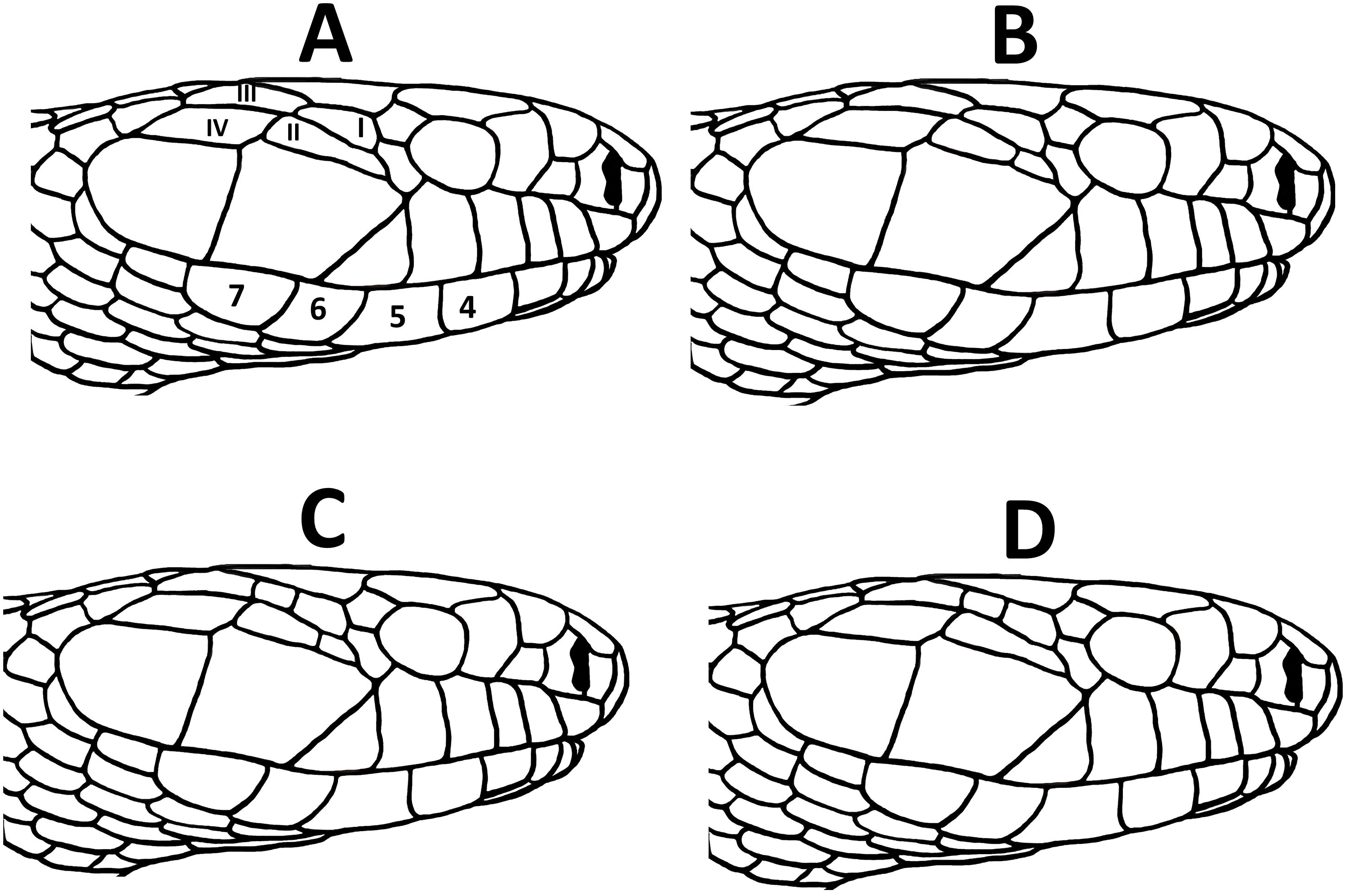
Head scale patterns in Eastern Indigo Snakes (*Drymarchon couperi*). A) 2+2 condition of temporals (I = dorsal anterior temporal; II = ventral anterior temporal; III = dorsal posterior temporal; IV = ventral posterior temporal) and position of 4^th^, 5^th^, 6^th^, and 7^th^ infralabials; B) 3_v_+2 condition of temporals (extra ventral temporal shaded); C) 4+2 condition of temporals (extra dorsal and ventral temporals shaded); D) 3_d_+2 condition of temporals (extra dorsal temporal shaded).

To better understand whether morphological variation provides secondary support for lineage divergence and speciation of *D. couperi*, we examined 114 individuals housed at the Orianne Center for Indigo Conservation, a sample that included 68 and 17 specimens from the Atlantic and Gulf lineages of Krysko et al. [15], respectively, along with 4 specimens of unknown lineage assignment, 12 specimens derived from F1 hybrids of Atlantic and Gulf lineage snakes, and 13 hybrids of Atlantic lineage snakes crossed with adults of unknown origin. Lateral or dorsolateral photos of the head were taken of each specimen, including a millimeter ruler for scale. The photos were used to record the condition of the temporal scales for each specimen. Four categories were recognized based on the number and position of temporal scales (Fig 1). We generated a contingency table providing counts of specimens in each of the four categories for each lineage, specimens of unknown lineage, and hybrids. Fisher’s exact test was used to determine whether the relative proportions of temporal scale categories differed between the Atlantic and Gulf lineages. Additionally, we measured total head length (posterior-most point of 8^th^ supralabial to anterior tip of rostral; N = 111), head height (only for photos in lateral aspect; at level of anterior-most point of parietal suture; N = 35), and length of the dorsal posterior temporal (intersection of ventral posterior temporal, dorsal posterior temporal, and adjacent first dorsal scale to intersection of ventral posterior temporal, dorsal posterior temporal, and adjacent ventral temporal; N = 111). All distances were measured using Adobe Photoshop 6.0 with reference to the photographed ruler. We used an analysis of covariance (ANCOVA) to test whether the linear relationship between head length and head height differed between Atlantic and Gulf lineages. We divided the length of the dorsal posterior temporal by head length to control for effects of body size and used an analysis of variance to test whether temporal length differed among the four categories of temporal scales.

We also examined 11 preserved specimens in the Auburn University Museum (AUM) collections. These snakes were from southeastern Georgia and presumed to belong to the Atlantic lineage. For these specimens, we measured length and width of the 6^th^ and 7^th^ infralabial scales with dial calipers. We measured both scales because it was not clear which of these was measured by Krysko et al. [16] and because they represent the position of the 7^th^ lower labial scale if the mental scale was included and if it was excluded. Additionally, we used photos of the type specimens presented in Krysko et al. [16] to determine length and width of the 6^th^ and 7^th^ infralabial scales using Adobe Photoshop 6.0. A length-to-width ratio was then calculated for each specimen. Mean differences between 6^th^ and 7^th^ infralabials were tested with a paired t-test. Differences between our sample of Atlantic lineage snakes and the type specimens was determined by visual inspection. We used SAS v.9.4 for all analyses (SAS Institute, Inc., 2008) with α = 0.05.

### Ethics Statement

The use of live snakes for research was approved by Auburn University IACUC protocols (PRN 2007-1142, 2010-1750, 2013-2386, 2017-3102) and a federal research permit from the United States Fish and Wildlife Service (TE32397A-O).

## Results

### Gene sequence analysis

We found that the nuclear locus NT3 was completely invariant across all *D. couperi* specimens, and thus NT3 had no variable sites for phylogenetic inference of *D. couperi*. As expected, the inferred ML phylogeny estimated a polytomy (Fig 2), indicating a lack of phylogenetic structure among individuals from the two mitochondrial lineages of *D. couperi* (*sensu lato*) [15]. Specimens of *D. melanurus erebennus* and *D. melanurus rubidus* clustered as sister taxa to the exclusion of all *D. couperi* specimens when rooted by *Coluber constrictor* + *Masticophis flagellum*.

**Fig 2.**
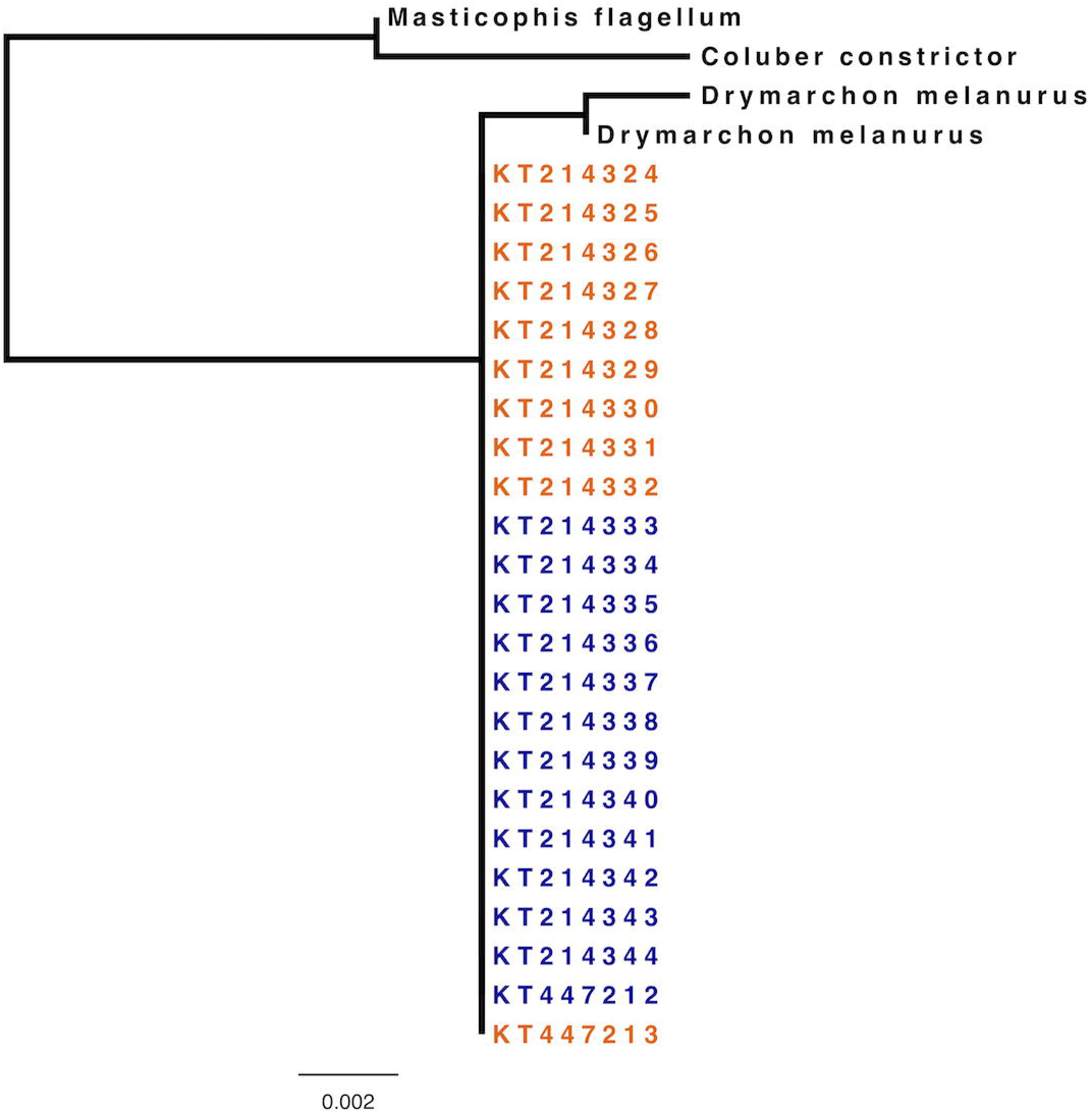
Maximum likelihood phylogeny among samples of Eastern Indigo Snakes (*Drymarchon couperi, sensu lato*) and outgroups (*Drymarchon melanurus*, *Coluber constrictor*, *Masticophis flagellum*), as inferred from sequence data from the nuclear gene *neurotrophin*-3 (NT3). Indigo snakes are labeled by GenBank accession numbers and are classified by the lineages identified by Krysko et al. [15]. Taxon labels: blue = Atlantic lineage, orange = Gulf lineage).

### Microsatellite analysis

We found evidence for null alleles in three loci (Dry33, Dry63, and Dry69). Therefore, we eliminated these loci from further analysis and retained the remaining 14 loci. Delta K was maximized at K = 2 (Fig 3) and supported two genetic clusters: one associated with the northernmost samples, and a second with the southern-most samples. However, there was extensive admixture between the two clusters, particularly in central Florida (Fig 4; Fig 5). For the 20 samples from Krysko et al. [15], including representatives from Atlantic and Gulf lineages, we found all individuals were assigned to our southern cluster, and all were highly admixed with the northern cluster (0.66 assignment to the southern cluster for the Gulf clade and 0.57 assignment to southern cluster for the Atlantic clade). Furthermore, the Jost’s D value for these 20 samples was extremely small (0.0004), indicating no genetic differentiation among samples at the putative contact zone. Raw likelihood values from Structure began to plateau at K = 3 (Fig 3), and we interpreted this as support for three populations: a cluster of the southern samples, but described further population subdivision of the northern samples into two clusters.

**Fig 3.**
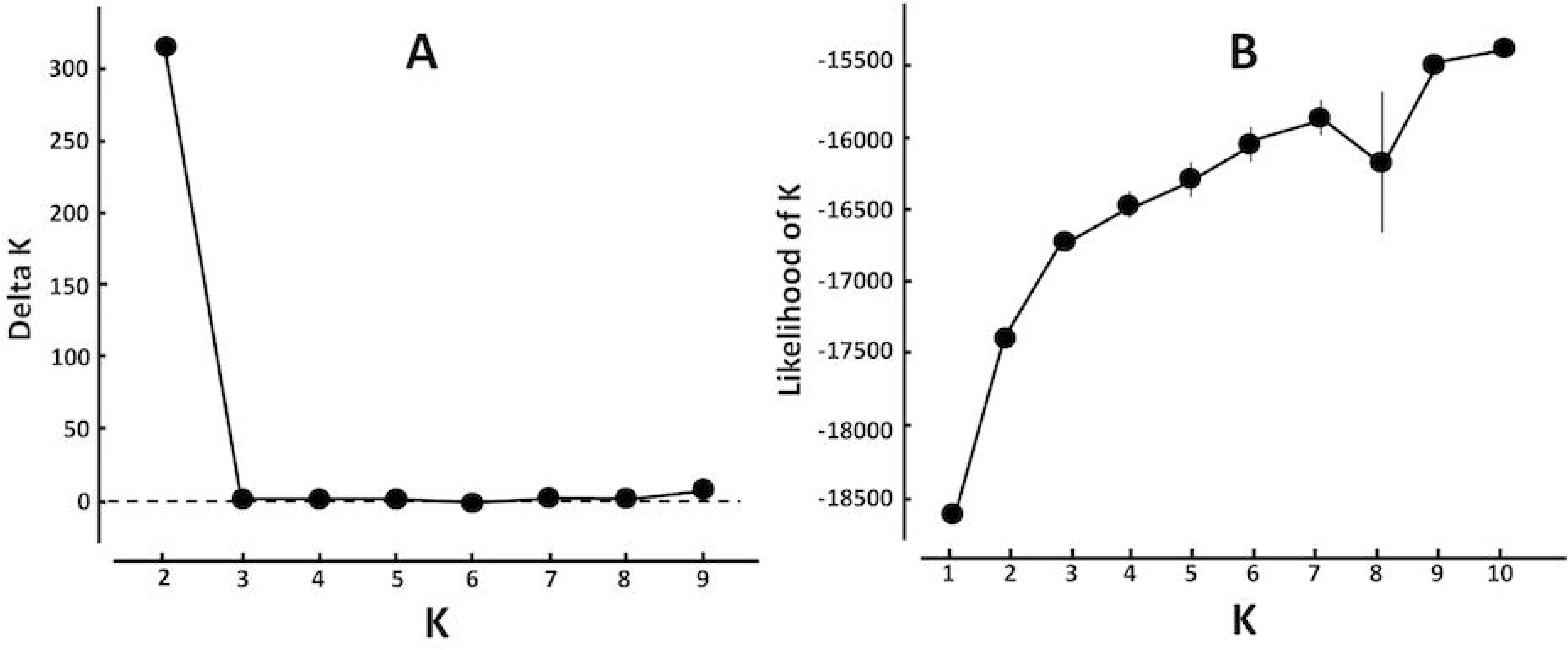
Plot of Delta K (A) and likelihood scores (B) used to identify the most likely number of population clusters across the range of *Drymarchon couperi* using the Bayesian algorithm Structure [43]. The dashed line in (A) indicates Delta K = 0; the error bars in (B) indicate S.D.

**Fig 4.**
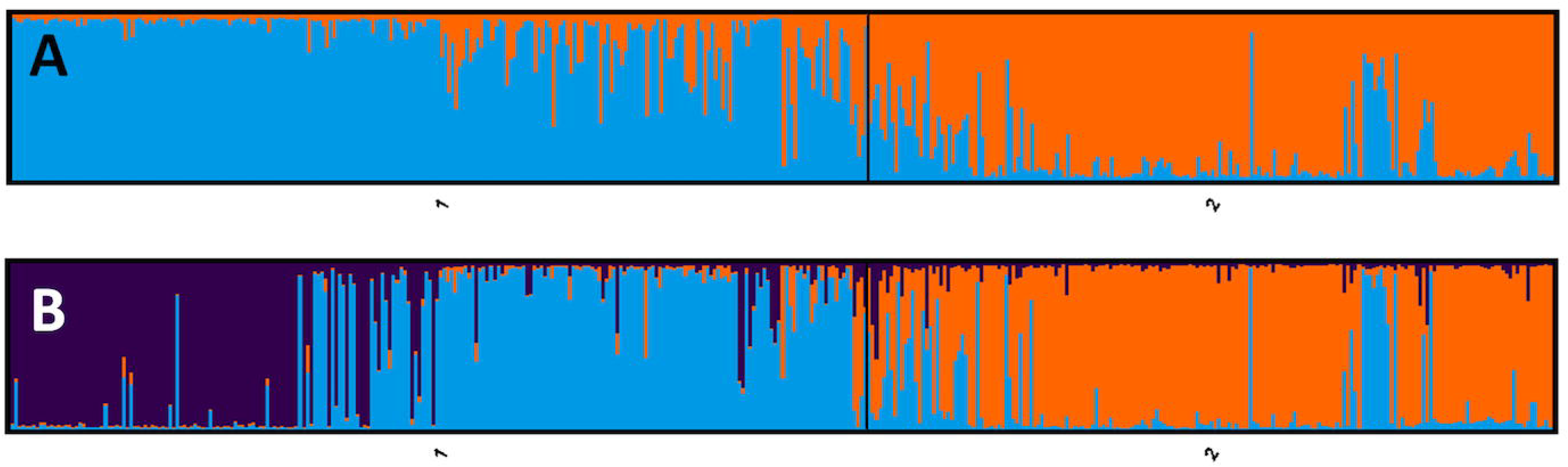
Bar plots of population clustering estimated through the Bayesian clustering algorithm Structure with (A) K = 2 and (B) K = 3. The y-axis is the proportion of individual ancestry for each cluster; in the x-axis, group 1 represents individuals assigned or assumed to be within the Atlantic mitochondrial clade, and group 2 represents individuals assigned or assumed to be in the Gulf mitochondrial clade. Within each group, individuals are sorted by latitude. (A) K = 2; blue indicates alleles from the northern population cluster, and orange indicates alleles from the southern population cluster. (B) K = 3; purple and blue indicate alleles from two northern population clusters, and orange indicates alleles from the southern population cluster.

**Fig 5.**
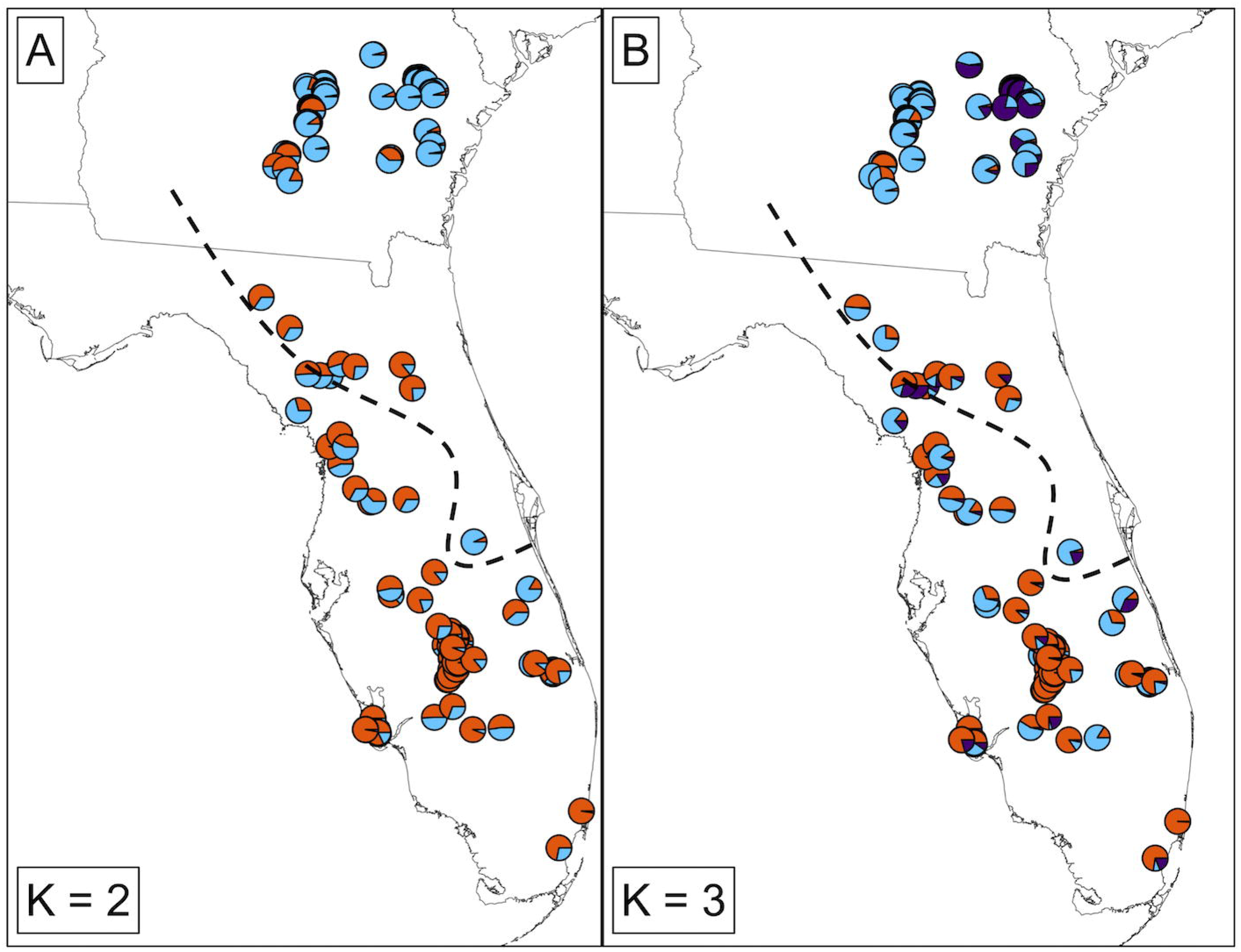
Maps of sampling sites represented as pie charts of percent ancestry within population clusters identified by Structure analyses. (A) K = 2 populations, with the southern cluster represented as black and the northern cluster as gray; (B) K = 3 populations, with the southern cluster as black and two northern populations as grey and white. Lines indicate state boundaries in the southeastern U.S.A. Pie charts for both panels are colored as in Fig 4.

Inferences about spatial genetic structure were very similar among the three connection networks we used, so we report the results using the distance-based connection network (see SI Figures S1 and S2 for results using the Delaunay triangulation network). Lagged scores from the first two PC axes were highly correlated between different connection networks (*r* ≥ 0.93). The global test was significant (observed = 0.0222, P < 0.0001) while the local test was not (observed = 0.0064, P = 0.84). The first PC axis explained the most variation followed by the second PC axis (0.2978 and 0.1533, respectively; all other axes ≤ 0.1002; SI Figure S3) and both axes showed positive spatial autocorrelation (Moran’s *I* = 0.83 and 0.74, respectively; SI Figure S3). The lagged scores of the first axis suggested genetic structure followed a north-south gradient, while the second axis suggested strongest genetic structure within southern Georgia and between Georgia and Florida (Fig 6). There was no consistent differentiation between samples from different lineages and relatively substantial overlap across the putative contact zone.

**Fig 6.**
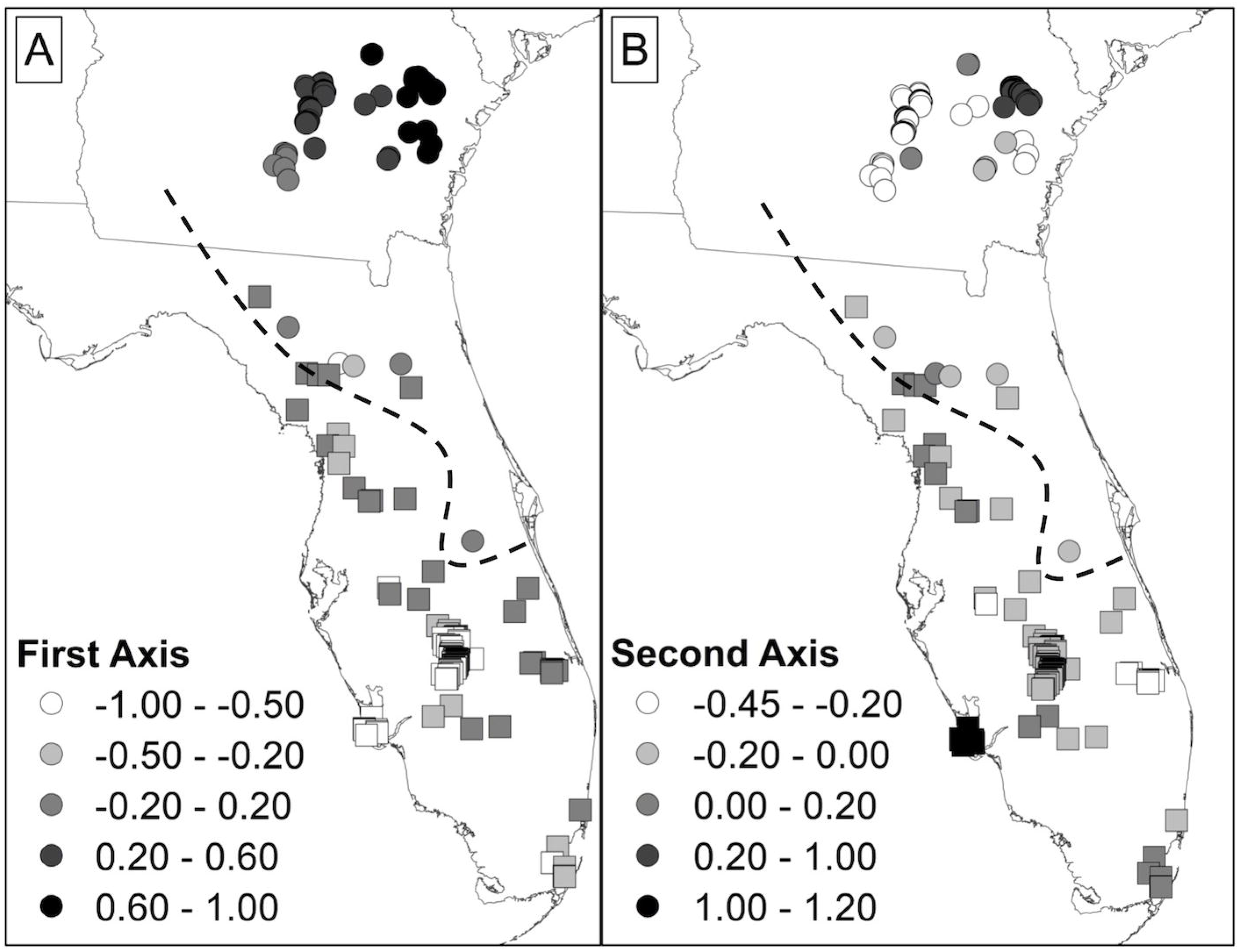
Spatially lagged scores for each sample from the first (A) and second (B) axes of a spatial principle components analysis (sPCA). Atlantic clade samples are displayed using circles while Gulf clade samples are displayed using squares. Samples with more extreme values/colors are more genetically differentiated.

MLPE models described strong effects of Euclidean distance on genetic distance among samples. Specifically, we observed significant effects of Euclidean distance on genetic distance within the Atlantic clade (β = 0.05, SE = 0.0007, *t* = 65.5, *R^2^* = 0.21), the Gulf clade (β = 0.07, SE = 0.001, *t* = 48.6, *R^2^* = 0.34), and among all samples (β = 0.04, SE = 0.0003, *t* = 148.8, *R^2^* = 0.18). When comparing genetic distance within and between mitochondrial clades, values within clades overlapped greatly with values observed among all samples between clades (Fig 7).

**Fig 7.**
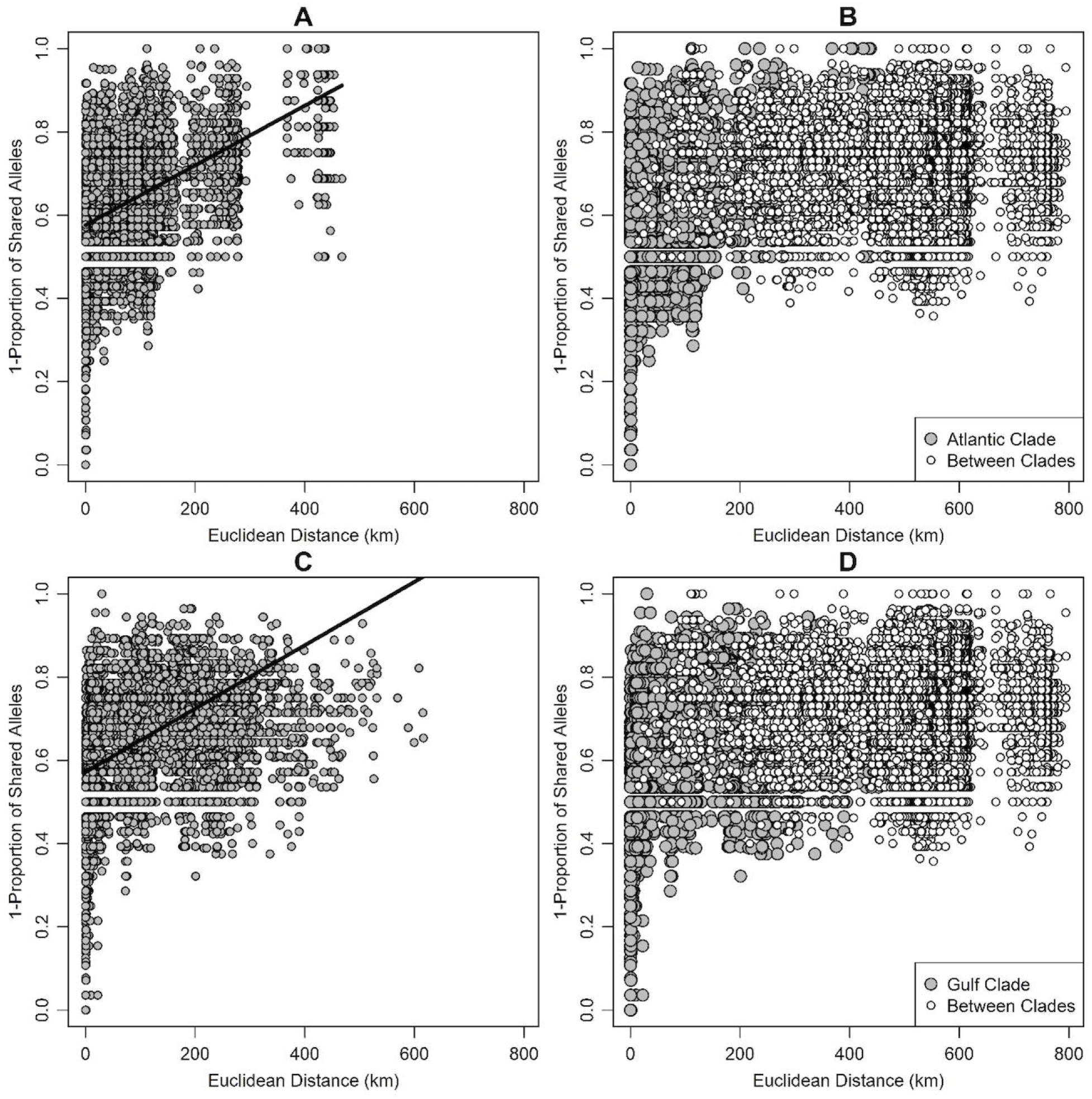
Plots of pairwise genetic distance (1 - proportion of shared alleles) against Euclidean distance (km) showing positive isolation by distance. Solid lines show the predicted pattern of isolation by distance from a linear mixed-effects model with maximum-likelihood population effects (MLPEs; [52]). (A) Pairwise distances within Atlantic clade samples; (B) pairwise distances within Atlantic clade samples (gray circles) and among samples from both Atlantic and Gulf clades (white circles); (C) pairwise distances within Gulf clade samples; (D) pairwise distances within Gulf clade samples (gray circles) and among samples from both Atlantic and Gulf clades (white circles).

### Morphological analysis

We observed four categories of temporal scales from both Atlantic and Gulf lineages (Table 1). In 22% of specimens, temporals conformed to the 2+2 formula that Krysko et al. [16] described as being invariant (Fig 1A). We found that 38% of specimens exhibited an extra ventral temporal (Fig 1B), 26% of specimens had extra dorsal and ventral temporals (Fig 1C), and 14% of specimens exhibited an extra dorsal temporal (Fig 1D). The frequency with which these four categories occurred differed between Atlantic and Gulf lineage specimens (Table 2; df = 3; Fisher’s Exact P = 0.006), with Atlantic lineage snakes tending to have two dorsal temporals and Gulf lineage snakes tending to have three dorsal temporals. Head shape, based on ANCOVA of head width on head length, did not differ between Atlantic and Gulf lineages in either slope (Fig 8; F = 0.07; df = 1; P = 0.79) or intercept (F = 0.48; df = 1; P = 0.49). Length of the dorsal posterior temporal, expressed as a proportion of head length, differed significantly among temporal categories (F = 18.34; df = 3; P < .0001), with the dorsal posterior temporal being proportionately shorter when three dorsal temporal scales are present relative to when two dorsal temporal scales are present (Fig 9).

**Table 1.** Frequency of occurrence of two and three dorsal temporal scales (DTs) between the Atlantic and Gulf lineages of Eastern Indigo Snakes.

| <u>Mitochondrial Clade</u> | <u>Scale Condition</u> |  |  |  |
| --- | --- | --- | --- | --- |
|  | 2+2 | 3<sub>v</sub>+2 | 3<sub>d</sub>+2 | 4+2 |
| Atlantic | 14 | 30 | 11 | 13 |
| Gulf | 5 | 1 | 2 | 9 |
| Atlantic x Gulf | 2 | 5 | 1 | 4 |
| Atlantic x Unknown | 2 | 6 | 1 | 4 |
| Unknown | 1 | 2 | 1 | 0 |

**Fig 8.**
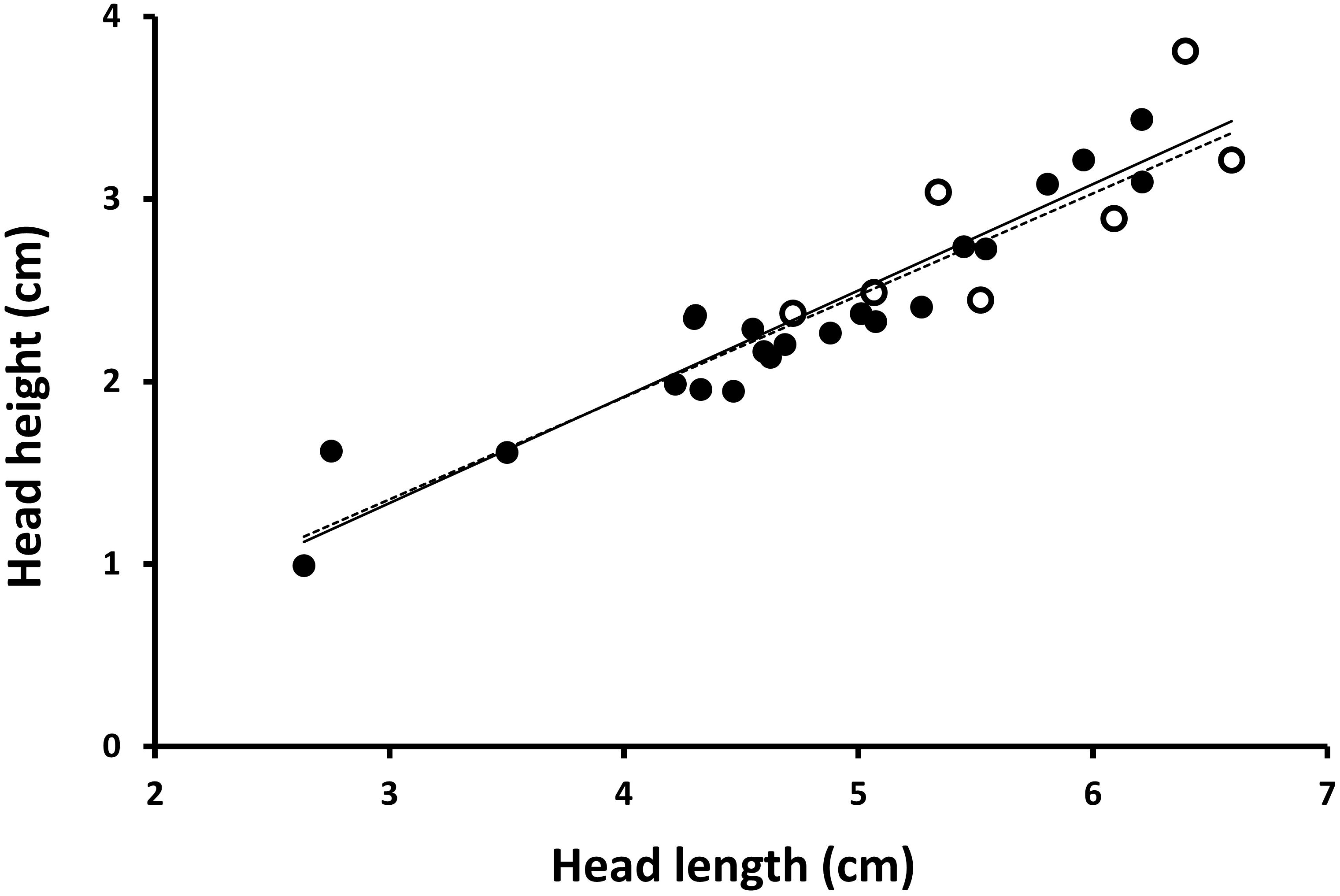
Bivariate plot of head height on head length. Values from Atlantic lineage indicated by solid circles and solid line; values from Gulf lineage indicated by open circles and dashed line.

**Fig 9.**
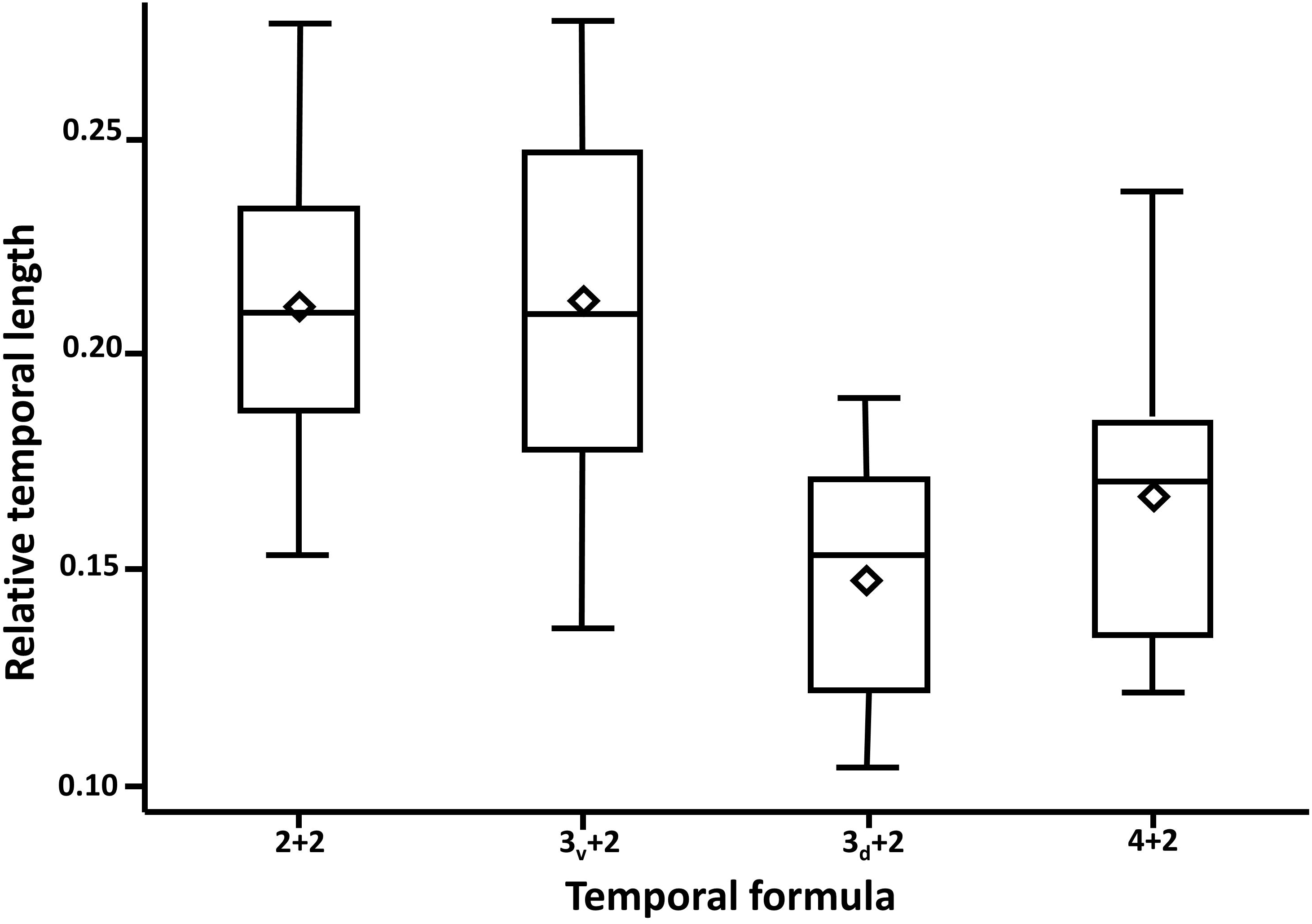
Box and whiskers plot of distribution of ratio of dorsal posterior temporal scale length to head length in four categories of temporal scales (see Fig 1). Vertical lines indicate range; box indicates interquartile, horizontal line indicates median; open diamond indicates mean.

When the length and width of the 6^th^ and 7^th^ infralabial scales were converted to a length-to-width ratio, the distribution of our sample of scales from Atlantic lineage snakes encompassed values for both type specimens for each scale (Fig 10). Length-to-width ratios differed between 6^th^ and 7^th^ infralabials (t = 8.07; df = 12, P < .0001), with 7^th^ infralabials being more elongate than 6^th^ infralabials.

**Fig 10.**
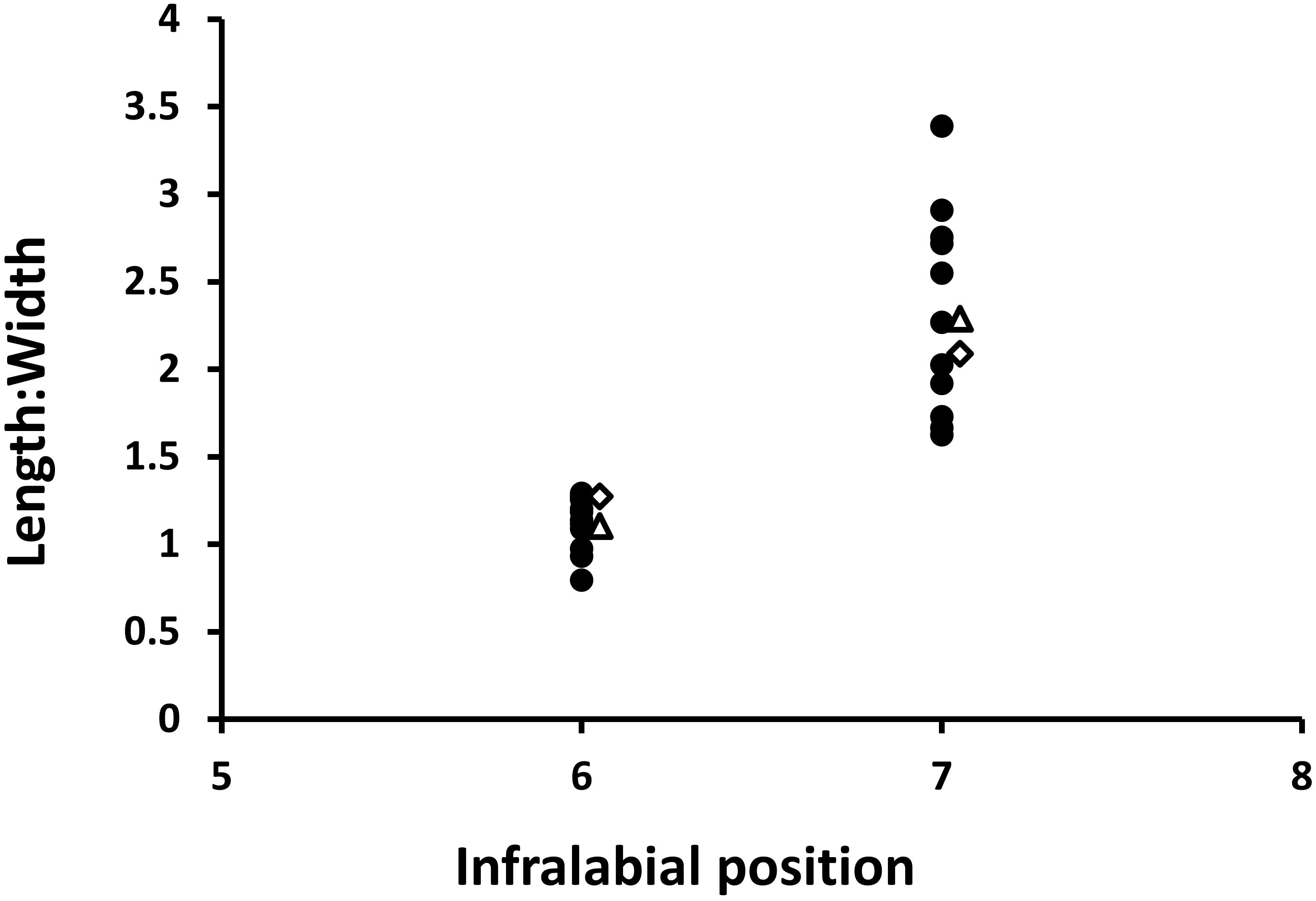
Distribution of length-to-width ratio of 6^th^ and 7^th^ infralabials in 11 Atlantic lineage specimens of *Drymarchon couperi* (dark spots). Open triangles indicate ratios from type specimen of *D. couperi* (Atlantic lineage); open diamonds indicate ratios from type specimen of *D. kolpobasileus* (Gulf lineage).

## Discussion

### Gene sequence analysis

We found no variation of the nuclear NT3 locus in sequences generated by Krysko et al. [15]. These authors included this locus because it “potentially represents an informative, single-copy, unlinked locus that is likely evolving at a different rate than mtDNA (mitochondrial) genes” (p. 114). From this we infer that the authors expected this nuclear gene to corroborate patterns generated from mtDNA, solidifying their conclusion that two species are present. However, if the locus was expected to be informative (i.e., *sensu* [57]), then the fact that it is invariant indicates that it (1) does not support population structure consistent with two genetic populations (i.e., two species), (2) was not necessary to include it in the study, and (3) only served to restrict phylogenetic inference to the mitochondrial genome and its inherent problems. From a statistical perspective, the only effect of including NT3 in their analysis was a slight reduction of branch lengths due to the sequence’s invariance. For these reasons, we suggest that the gene sequence data do not provide compelling evidence in support of two distinct species of *D. couperi* (*sensu lato*).

### Microsatellite analysis

Our population genetic analyses found evidence of population structure within Eastern Indigo Snakes, particularly along a north-south gradient and between southern Georgia and peninsular Florida. However, patterns of microsatellite genetic structure reveal substantial differences from the phylogenetic structure inferred from Krysko et al. [15]. First, the geographic pattern of our population clusters suggests a north-south orientation rather than the Gulf-Atlantic orientation suggested by mtDNA. Second, and most importantly, our Structure and sPCA analyses document widespread contemporary admixture of alleles from the northern and southern populations. Admixture occurs across the entire range of *D. couperi* (*sensu lato*) and cannot be characterized as a narrow hybrid zone. While our MLPE analyses found support for isolation by distance both across all samples and within each mtDNA clade, those analyses also found high degree of overlap in genetic distance within and between clades, an observation that we view as inconsistent with a two-species hypothesis. Further, our sPCA results, which accounted for spatial autocorrelation among samples, are consistent with isolation by distance along a north-south gradient and are not consistent with the statement by Krysko et al. [15] that the two lineages of have “near discrete geographic distributions” (p. 118) and that “dispersal between lineages is too low to influence demographic processes” (p. 119). Rather, our results indicate high levels of contemporary gene flow, and we instead suggest that genetic structure among populations is best described as continuous isolation by distance rather than discrete evolutionary lineages. While isolation by distance is possible across two discrete clusters [58], multiple analyses of genetic variation within our microsatellite data, including our sPCA which accounts for spatial autocorrelation among samples, nevertheless did not correspond to the genetic lineages recognized by Krysko et al. [15,16]. Together, we interpret the microsatellite results as failure to support the presence of two discrete evolutionary metapopulations with little or no admixture, the necessary criterion for the existence of two species [20]. Thus, we view these data as sufficient to reject the two-species hypothesis, and we suggest the available data are most consistent with the existence of a single evolutionary species, *D. couperi*.

The North American Coastal Plain is a global biodiversity hotspot [59]. In particular, peninsular Florida has a high proportion of endemic species (e.g., snakes: [60–62]), which are likely a product of refugial isolation on islands during periods of elevated sea level in recent epochs and contemporary drainage-driven endemism [63,64]. However, our case study with *D. couperi* adds to a growing number of examples of Florida organisms that appear, based on modeling of one or few genetic loci, to represent species that are distinct from other mainland counterparts, but for which microsatellite or similar data demonstrate substantial gene flow. Burbrink and Guiher [60] estimated that there was such low gene flow between cottonmouths (*Agkistrodon piscivorus*) in Florida and the mainland that speciation must have occurred between those two regions, a hypothesis immediately contested by data from Strickland et al. [65] who detected a broad geographic range of admixture using AFLP markers. Similarly, Thomas et al. [66] described alligator snapping turtles (*Macrochelys temminckii*) from the Apalachicola River and adjacent rivers to be a distinct species, despite microsatellite data from Echelle et al. [67] that are inconsistent with this conclusion [68]. In a similar example, Gordon et al. [69] used mtDNA to describe a population of *Anaxyrus boreas* in the western United States as a distinct species, but this conclusion was contested by microsatellite analysis of gene flow [70].

We believe this emerging pattern along with a concept of species as evolutionary metapopulation lineages [20] can guide us toward a more rigorous framework for species delimitation. While analyses of one or few genetic loci can be informative by revealing apparent monophyly, they should not be viewed as sufficient to diagnose and delimit species, because of high error rate associated with the gene tree/species tree problem [24,25]. This issue is particularly acute when relying on mtDNA [4,26]. We suggest that authors be required to meet the necessary criterion of the unified species concept – that species are independently evolving metapopulations lineages – by demonstration of little or no contemporary gene flow across lineage populations using a multi-locus or genomic dataset. We note that Krysko et al. [16] also used the unified species concept [20] as a framework to define species; while details about how they operationally diagnosed species were not clear, it appears that they relied on evaluation of reciprocal monophyly of mtDNA. While monophyly can be used as secondary criterion under the metapopulation lineage concept [20], for the reasons outlined above we suggest authors should evaluate monophyly in conjunction with analyses of contemporary gene flow and population admixture, particularly when monophyly is evaluated with small numbers of loci.

### Morphological analysis

Contrary to the results presented in Krysko et al. [16], we rejected the hypothesis that the Atlantic and Gulf lineages are identifiable entities revealed by morphology. We reached this conclusion after re-examining the variables used by Krysko et al. [16] to diagnose each lineage. Of the disparities that emerge between our analyses and theirs, the conformation of the infralabials is the most problematic. The figures presented by Krysko et al. [16] for the 6^th^ infralabial show great promise for diagnosing lineages, and separation of the lineages along PC1 of their analysis seems to provide statistical support for this character. However, we were struck by how dissimilar Atlantic specimens appeared to be from the scale shape ascribed to them by Krysko et al. [16]. Our analyses demonstrate that the 6^th^ and 7^th^ infralabials differ in shape, that the shape of the 7^th^ infralabial conforms to the shape ascribed to the Atlantic lineage, and that the shape of the 6^th^ infralabial conforms to that ascribed to the Gulf lineage. It is unclear which of these scales was measured by Krysko et al. [16] and we found that the range of variation of each scale within a sample of Atlantic lineage snakes encompasses both type specimens. One potential explanation for this discrepancy is that Krysko et al. [16] intended to measure the 7^th^ infralabial but inadvertently measured the 6^th^ for Gulf lineage specimens and the 7^th^ for Atlantic lineage specimens, perhaps because the mental scale sometimes was included in counts and other times was not. Otherwise, we are left with a PCA that separates lineages based on one of the two possible scales, but a univariate analysis that fails to confirm these lineages.

Our results for the temporal scale reveal great variation in the number of these scales present in Eastern Indigo Snakes. The four categories that characterize this variation are found in both Atlantic and Gulf lineage snakes, indicating that this feature is not diagnostic. Nevertheless, Atlantic lineage snakes tend to have two dorsal temporals, while Gulf lineage snakes tend to have three. We assume that Krysko et al. [16] intended to measure the dorsal posterior temporal and, therefore, we focused our attention on this scale. Our data indicate that the length of the dorsal posterior temporal, relative to head length, becomes shortened if three dorsal temporals are present and becomes elongate if two dorsal temporals are present. This finding indicates that the scale shapes revealed by PC2 of Krysko et al. [16] represent distinguishable groups, but these represent two phenotypes and not two species. We infer that the different morphologies of the dorsal posterior temporal result because, during embryonic development of some individuals, the dorsal anterior temporal divides, limiting space for development of the dorsal posterior temporal.

Krysko et al. [16] also used head shape to diagnose the two lineages. Relative head length and head height did load heavily on their PC1 axis and they used this to diagnose the Atlantic lineage as having an elongate wide head and to diagnose the Gulf lineage as having a short narrow head. Our bivariate examination of head length and height revealed no difference in head shape between the two lineages. We are unsure why our results differed from Krysko et al. [16], although we note that snake morphology can be difficult to measure consistently [71]. Specimens preserved with mouths open are likely to have larger values for head height than those with mouths closed. If the relative frequency of open-mouthed versus closed-mouthed specimens differed between lineages, this might yield a spurious association of head shape with lineage. Our measurements were made from live specimens with closed mouths, which we assume reduces measurement error. If the lineages truly differ in head shape, our ANCOVA should have revealed this difference.

### Natural history considerations

Our examination of genetic and morphological variation in *D. couperi* (*sensu lato*) demonstrates that the two-species hypothesis proposed by Krysko et al. [16] is not supported by available data. We offer several explanations for why true patterns of gene flow might result in a lack of genetic and morphological differentiation. First, *D. couperi* movements can be extensive, especially for males. Male annual home range size is as large as ca. 1500 ha [72] and average ca. 2.5–6.6 times larger than for females [72,73]. In fact, the disparity between male and female home range sizes becomes exacerbated in large snakes, a feature dominated by data from *D. couperi* (Fig 11; S3 Table). Within peninsular Florida, male *D. couperi* can move up to ca. 2 km in a single day and average movement distance in males is approximately twice that of females [38, D. Breininger, unpublished data]. Furthermore, males within peninsular Florida increase their movement frequency, distance, and home range size during the breeding season [38,74]. Dispersal distance of males may be 10 times that of females [75], with a small adult male in southern Georgia dispersing at least 22.2 km (straight line) over approximately two years [50].

**Fig 11.**
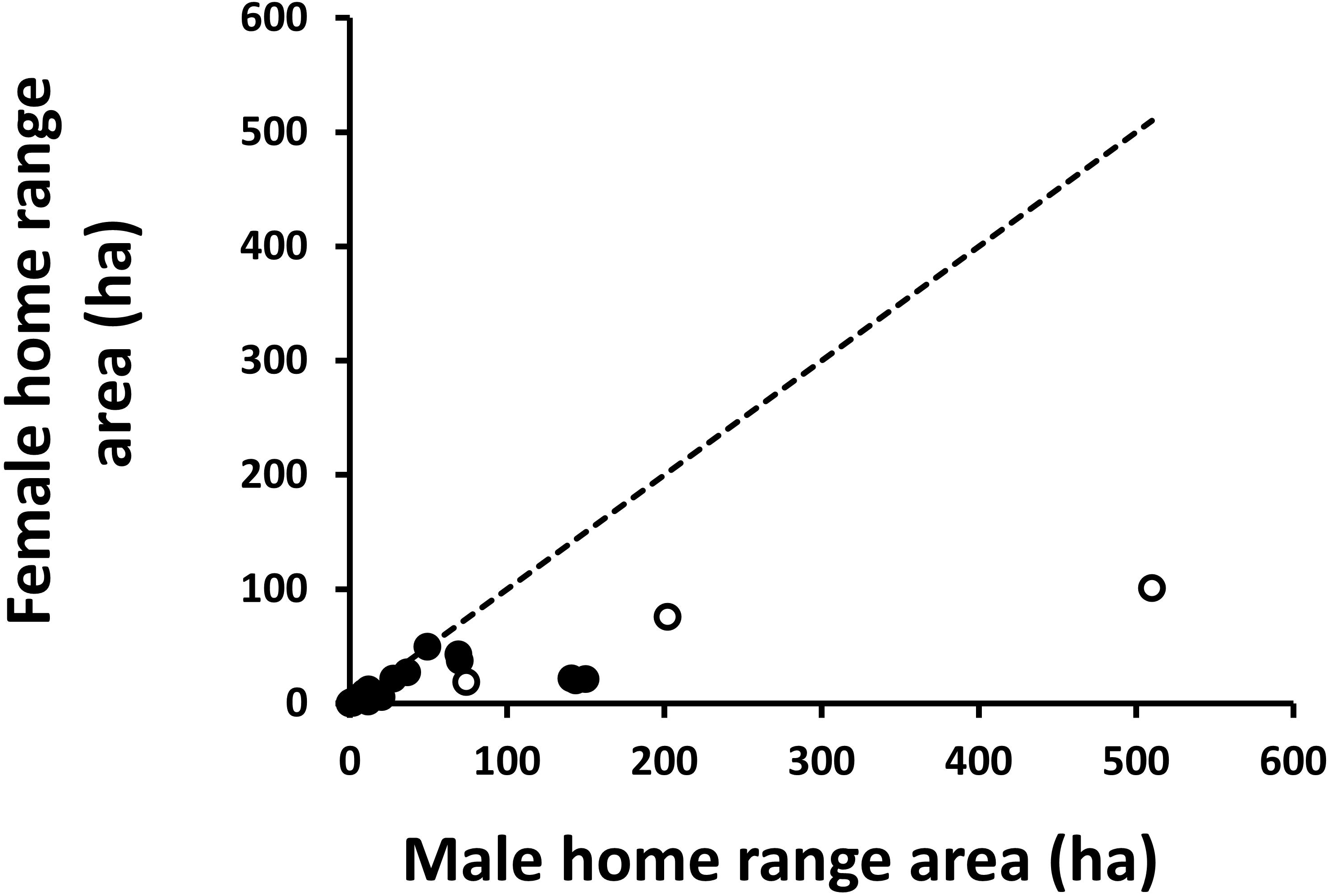
Bivariate plot of female home range area (ha) on male home range area for 25 studies of 21 species of snakes (S3 Table). Open circles are data from *Drymarchon couperi*; closed circles are data from all other species of snakes. Dashed line is null expectation if male and female home range sizes are equal in size.

A second feature of *D. couperi* life history that reduces opportunity for speciation is the variety of habitats used by these animals throughout a year (e.g., [72]), particularly in peninsular Florida, where individuals will utilize habitats with varying degrees of anthropogenic disturbance [76]. This broad habitat use reduces the opportunity for ecological barriers to gene flow and is consistent with the high rates of gene flow indicated by our microsatellite analysis. Additional life history observations show that *D. couperi* (*sensu lato*) can cross fresh and saltwater features 6–264 m wide [77; D. Stevenson, personal observation; D. Breininger, unpublished data], and we note that traditional river barriers [63] do not appear to limit gene flow in these large snakes. While we do not doubt that a historical climatic event may have separated *D. couperi* into two populations [15], the observed levels of contemporary gene flow indicate that genetic populations of *D. couperi* are in the process of merging back into a single evolutionary population (i.e., species), as has been observed for diverse taxa following climatic cycles [78].

Together, our knowledge of *D. couperi* movement patterns and life history allows us to hypothesize that limited female movement drives structure of the maternally-inherited mtDNA data presented by Krysko et al. [15], even though high levels of male movement drive extensive gene flow of the nuclear genome [79]. This life history-based model of intersexual variance in *D. couperi* gene flow is consistent with two other reptile systems for which life-history strategies generate contrasting patterns of gene flow among populations. First, female philopatry of Loggerhead Sea Turtles (*Caretta caretta*) causes structuring of mtDNA in the Atlantic Ocean, but high male dispersal drives significant nuclear gene flow among populations [80]. Second, a recent study of Neotropical snakes found active-foraging species to have greater rates of nuclear gene flow than ambush-predator species with more limited dispersal [81]. Last, and most relevant, many squamates are polygynous and are characterized by greater male dispersal relative to females [Fig 11; 82–85]; therefore, squamates typically have greater nuclear gene flow than that of the mitochondrion [79]. This pattern is also seen in waterfowl [28,30]. Thus, we suggest that *D. couperi* is similar to other reptile species in that life history provides explanations for why phylogeographic patterns from mtDNA are inconsistent with historical and/or contemporary patterns of nuclear DNA. Because of this incongruence and the high levels of recent gene flow among populations observed by our microsatellite analysis, we place *Drymarchon kolpobasileus* into synonomy with *D. couperi*. For now, we retain all Eastern Indigo Snakes within *D. couperi*, but note that the analysis of Krysko et al. [16] reveals that this species may not be diagnosable from *D. melanurus erebennus*.

### Conservation implications

Eastern Indigo Snakes are now being released into the Conecuh National Forest of south-central Alabama and to the Apalachicola Bluffs and Ravines Reserve in the panhandle of Florida. Efforts to repatriate the species are the result of extensive collaborations and partnerships between state and federal agencies, as well as non-governmental conservation organizations. Initially, snakes used for release were to come from wild-caught gravid females taken from sites in southeastern Georgia, retained in captivity until they laid eggs, and then released at the point of capture. Eggs from these initial females then were to be hatched in captivity and raised for release at one of the two repatriation sites or donated to the Orianne Center for Indigo Conservation, where captive breeding from these and other donated animals was to occur, eventually replacing use of wild-captured females.

Those charged with implementing conservation planning for *D. couperi* welcome the opportunity to base those plans on the best published science. Many meetings were held among stakeholders and scientists, and, as Krysko et al. [15] noted, their preliminary molecular results [86] were presented at one of these early meetings as a challenge to the release of Atlantic lineage snakes at the two release sites. Here we describe why data from that preliminary report did not compel changes in the repatriation plan, and why continuation of that plan is unlikely to encounter the problems predicted by Krysko et al. [15].

A lack of apparently viable populations of Eastern Indigo Snakes in the Florida panhandle and southern Alabama and Mississippi was recognized as a significant gap in efforts to retain Eastern Indigo Snakes across its historic geographic range. This gap guided the choice of the two sites where repatriation was deemed reasonable [13]. Data from Krysko et al. [86] were used to propose that only snakes of the Gulf lineage be used for repatriation because genetic information from two isolated samples from the panhandle of Florida were of this lineage. However, because Eastern Indigo Snakes were extirpated at both sites, discussions centered on how to deal with the uncertainty of which lineage historically occurred at each site and, therefore, which lineage(s) were appropriate to release at each site. This uncertainty recognized the fact that, for both sites, the 3^rd^ and 4^th^ closest localities currently occupied by Eastern Indigo Snakes have the Atlantic lineage. Additionally, the Atlantic lineage occurs in the Suwannee drainage, which empties into the Gulf of Mexico, further illustrating the uncertainty in identifying the location of the Gulf-Atlantic split and that both lineages are found in north Florida where they might serve as colonists for the release sites. Participants in these discussions were made aware of then-unpublished microsatellite data demonstrating widespread gene flow between lineages and observations from the zoo and herpetocultural communities documenting interbreeding of the two lineages with no apparent effects of outbreeding depression. While there is currently no evidence of ecological differences between Atlantic and Gulf lineage snakes, snakes from northern populations are known to differ markedly from southern populations in seasonal activity. Specifically, *D. couperi* in the latter region do not appear dependent upon Gopher Tortoise (*Gopherus polyphemus* Daudin 1802) burrows for winter refugia, likely because of milder winter temperatures [38,72,73,87,88]. As a result, use of snakes from similar latitudes appeared to be more important in ensuring successful repatriation than use of snakes from the same mitochondrial lineage. The majority of meeting participants therefore concluded that the wide movements of male *D. couperi* and broad habitat use of the species made matching of mitochondrial DNA lineages less important in guiding repatriation than matching spatial variation in ecology. Preservation of mitochondrial lineages was already achieved through apparently viable populations at the northern and southern geographic extremes of *D. couperi* and release of snakes of one mitochondrial lineage into an area historically occupied by the other lineage was viewed simply as replicating patterns of gene flow observed throughout much of north and central Florida.

Because inferences from phylogenetic analyses of few loci can conflict with microsatellite-based analyses of contemporary admixture, we suggest that, at a minimum, both tools should be considered to adequately reveal recently diverged taxa. Additionally, we argue that a more-consistent voice from the community of systematists will emerge when life-history data are allowed to inform speciation models. For example, the potential for large snakes to move long distances as well as a difference in movement behavior between the sexes limit the utility of mitochondrial data alone to reveal novel species. We suspect this disparity in movement is rampant within squamates (e.g., [89]). Finally, species descriptions for which diagnoses merely report measures of variables that overlap widely among taxa (e.g., [16,66]) or that are not derived from results presented in the paper (e.g., [60]) should be rejected because they fail to provide further evidence that proposed taxa are identifiable as individuals. We point to papers that carefully meld phylogenetic and population genetic analyses [70,90,91] as improved examples of the process by which researchers might determine whether several populations have diverged sufficiently to be recognized as distinct species. Last, we suggest authors and reviewers be particularly critical of species descriptions without careful analysis of admixture or contemporary gene flow, because these papers may incorrectly delimit species, contribute to erroneous hyperdiversity, and confuse efforts to understand and conserve imperiled biodiversity.

## Acknowledgments

We thank the Alabama Department of Conservation and Natural Resources, Auburn University College of Science and Mathematics, Orianne Society, and U.S. Fish and Wildlife Service for funding that made this study possible. We thank the Orianne Center for Indigo Conservation for allowing us access to the captive breeding collection of *D. couperi*, and we thank the Florida Museum of Natural History, M. Aldecoa, P. Barnhart, D. Ceilley, Z. Forsburg, G. Graziani, C. Lechowicz, J. Macey, S. Mortellaro, L. Paden, D. Parker, K. Ravenscroft, M. Ravenscroft, B. Rothermel, A. Safer, J. Sage, F. Snow, K. Stohlgren, M. Wallace, Sr., and J. Wasilewski for providing samples. Our manuscript benefitted greatly from comments made by anonymous reviewers. This paper is contribution no. […] of the Auburn University Museum of Natural History.

## Author Contributions

**Conceived and designed the experiments:** BF JB SS CG.

**Project coordination:** BF.

**Contributed reagents/materials/analysis tools:** JB SS CJ CG.

**Collected the data:** BF JB SP MH CG.

**Analyzed the data:** BF JB SS CG.

**Wrote the paper:** BF JB SS DS JRO DAS CG.

## Supporting Information

S1 Fig. Results of spatial principle components analysis using a Delaunay triangulation connection network. The eigenvalues (left) for each axis where positive values indicate global structure and negative values indicate local structure, and the Moran’s *I* (right) plotted against the variance for each axis.

S2 Fig. Additional results of spatial principle components analysis using a Delaunay triangulation connection network demonstrating the spatially lagged scores from the first (A) and second (B) axes. Atlantic clade samples are displayed using circles while Gulf clade samples are displayed using squares. Samples with more extreme values/colors are more genetically differentiated.

S3 Fig. Results of spatial principle components analysis using a distance-based connection network where samples are considered connected if they are within 22.2 km which is the maximum recorded dispersal distance for Eastern Indigo Snakes. The left figure shows the eigenvalues for each axis where positive values indicate global structure and negative values indicate local structure and the right figure shows the Moran’s *I* plotted against the variance for each axis.

S1 Table. GenBank accession numbers for sequence data from the nuclear gene *neurotrophin*-3 (NT3) from Krysko et al. [15] that were analyzed to generate Fig 2.

S2 Table. Multiplex PCR panels for *Drymarchon couperi* microsatellite loci. The names of loci are as in [39].

S3 Table. Citations for snake studies (N = 28) describing intersexual variance in home range size (hectares) for 22 species of snakes used to generate Fig 9. See [92] for an earlier review of the topic.

## References

1. Avise JC. A role for molecular genetics in the recognition and conservation of endangered species. Trends Ecol Evol. 1989;4: 279–281.

2. Agapow P-M, Bininda-Emonds OR, Crandall KA, Gittleman JL, Mace GM, Marshall JC, et al. The impact of species concept on biodiversity studies. Q Rev Biol. 2004;79: 161– 179. doi:10.1086/383542

3. Mace GM. The role of taxonomy in species conservation. Philos Trans R Soc Lond B Biol Sci. 2004;359: 711–9. doi:10.1098/rstb.2003.1454

4. Frankham R, Ballou JD, Dudash MR, Eldridge MDB, Fenster CB, Lacy RC, et al. Implications of different species concepts for conserving biodiversity. Biol Conserv. Elsevier Ltd; 2012;153: 25–31. doi:10.1016/j.biocon.2012.04.034

5. Daugherty C, Cree A, Hay J, Thompson M. Neglected taxonomy and continuing extinctions of tuatara (*Sphenodon*). Nature. 1990;374: 177–179. doi:10.1016/0021-9797(80)90501-9

6. Ghiselin M. Metaphysics and the origin of species. SUNY Press, editor. Albany, New York; 1987.

7. Carstens BC, Pelletier TA, Reid NM, Satler JD. How to fail at species delimitation. Mol Ecol. 2013;22: 4369–83. doi:10.1111/mec.12413

8. US Fish and Wildlife Service. Eastern indigo snake recovery plan. Atlanta, Georgia; 1982.

9. US Fish and Wildlife Service. Eastern Indigo Snake (Drymarchon couperi): 5-year Review-Summary and Evaluation. Jackson, MS; 2008.

10. US Fish and Wildlife Service. Endangered and Threatened Wildlife and Plants. Listing of the Eastern indigo snake as a threatened species. Fed Regist. 1978;43: 4026–4029.

11. Stevenson DJ, Enge KM, Carlile LD, Dyer KJ, Norton TM, Hyslop NL, et al. An Eastern Indigo Snake (*Drymarchon couperi*) mark-recapture study in southeastern Georgia. Herpetol Conserv Biol. 2009;4: 30–42.

12. Hyslop NL, Stevenson DJ, Macey JN, Carlile LD, Jenkins CL, Hostetler JA, et al. Survival and population growth of a long-lived threatened snake species, *Drymarchon couperi* (Eastern Indigo Snake). Popul Ecol. 2012;54: 145–156. doi:10.1007/s10144-011-0292-3

13. Enge KM, Stevenson DJ, Elliott MJ, Bauder JM. The historical and current distribution of the Eastern Indigo Snake (*Drymarchon couperi*). Herpetol Conserv Biol. 2013;8: 288– 307.

14. Breininger DR, Legare ML, Smith RB. Eastern Indigo Snakes (*Drymarchon couperi*) in Florida: Influence of Edge Effects on Population Viability. Ecosystems. 2004.

15. Krysko KL, Nuñez LP, Lippi CA, Smith DJ, Granatosky MC. Pliocene-Pleistocene lineage diversifications in the Eastern Indigo Snake (*Drymarchon couperi*) in the Southeastern United States. Mol Phylogenet Evol. Elsevier Inc.; 2016;98: 111–122. doi:10.1016/j.ympev.2015.12.022

16. Krysko KL, Granatosky MC, Nuñez LP, Smith DJ. A cryptic new species of Indigo Snake (genus *Drymarchon*) from the Florida Platform of the United States. Zootaxa. 2016;4138: 549–569.

17. Pauly GB, Piskurek O, Shaffer HB. Phylogeographic concordance in the southeastern United States: the flatwoods salamander, *Ambystoma cingulatum*, as a test case. Mol Ecol. 2007;16: 415–29. doi:10.1111/j.1365-294X.2006.03149.x

18. Soltis PS, Gitzendanner MA. Molecular Systematics and the Conservation of Rare Species. Conserv Biol. 1999;13: 471–483.

19. de Queiroz K. Ernst Mayr and the modern concept of species. Proc Natl Acad Sci U S A. 2005;102 Suppl: 6600–7. doi:10.1073/pnas.0502030102

20. de Queiroz K. Species Concepts and Species Delimitation. Syst Biol. 2007;56: 879–886. doi:10.1080/10635150701701083

21. Will KW, Mishler BD, Wheeler QD. The Perils of DNA Barcoding and the need for integrative taxonomy. Syst Biol. 2005;54: 844–851.

22. Dayrat B. Towards integrative taxonomy. Biol J Linn Soc. 2005;85: 407–415. doi:doi:10.1111/j.1095-8312.2005.00503.x

23. Funk DJ. Molecular systematics of cytochrome oxidase I and 16S from *Neochlamisus* leaf beetles and the importance of sampling. Mol Biol Evol. 1999;16: 67–82.

24. Funk DJ, Omland KE. Species-level paraphyly and polyphyly: frequency, causes, and consequences, with insights from animal mitochondrial DNA. Annu Rev Ecol Evol Syst. 2003;34: 397–423. doi:10.1146/annurev.ecolsys.34.011802.132421

25. Dupuis JR, Roe AD, Sperling FAH. Multi-locus species delimitation in closely related animals and fungi: one marker is not enough. Mol Ecol. 2012;21: 4422–4436. doi:10.1111/j.1365-294X.2012.05642.x

26. Ballard JWO, Whitlock MC. The incomplete natural history of mitochondria. Mol Ecol. 2004;13: 729–744. doi:10.1046/j.1365-294X.2003.02063.x

27. Irwin DE. Phylogeographic breaks without geographic barriers to gene flow. Evolution (NY). 2002;56: 2383–2394. doi:10.1554/0014-3820(2002)056[2383:pbwgbt]2.0.co;2

28. Scribner KT, Petersen MR, Fields RL, Talbot SL, Pearce JM, Chesser RK, et al. Sex-Biased Gene Flow in Spectacled Eiders (Anatidae): Inferences from Molecular Markers with Contrasting Modes of Inheritance. Evolution (NY). 2001;55: 2105–2115.

29. Johansson H, Surget-Groba Y, Thorpe RS. Microsatellite data show evidence for male-biased dispersal in the Caribbean lizard *Anolis roquet*. Mol Ecol. 2008;17: 4425–4432. doi:10.1111/j.1365-294X.2008.03923.x

30. Peters JL, Bolender KA, Pearce JM. Behavioural vs. molecular sources of conflict between nuclear and mitochondrial DNA: The role of male-biased dispersal in a Holarctic sea duck. Mol Ecol. 2012;21: 3562–3575. doi:10.1111/j.1365-294X.2012.05612.x

31. Gamble T, Berendzen PB, Bradley Shaffer H, Starkey DE, Simons AM. Species limits and phylogeography of North American cricket frogs (*Acris*: Hylidae). Mol Phylogenet Evol. 2008;48: 112–25. doi:10.1016/j.ympev.2008.03.015

32. Grismer JL, Bauer AM, Grismer LL, Thirakhupt K, Aowphol A, Oaks JR, et al. Multiple origins of parthenogenesis, and a revised species phylogeny for the Southeast Asian butterfly lizards, *Leiolepis*. Biol J Linn Soc. 2014;113: 1080–1093. doi:10.1111/bij.12367

33. Folt B, Garrison N, Guyer C, Rodriguez J, Bond JE. Phylogeography and evolution of the Red Salamander (*Pseudotriton ruber*). Mol Phylogenet Evol. Elsevier Inc.; 2016;98: 97– 110. doi:10.1016/j.ympev.2016.01.016

34. Kimura M. A simple method for estimating evolutionary rate of base substitutions through comparative studies of nucleotide sequences. J Mol Evol. 1980;16: 111–120.

35. Schliep K. phangorn: phylogenetic analysis in R. Bioinformatics 27: 592–593.

36. R Core Team. R: A language and environment for statistical computing. In: R Foundation for Statistical Computing, Vienna, Austria [Internet]. 2016. Available: https://www.rproject.org

37. Bauder JM, Barnhart P. Factors affecting the accuracy and precision of triangulted radio telemetry locaitons of Eastern Indigo Snakes. Herpetol Rev. 2014;45: 590–597.

38. Bauder J, Breininger DR, Bolt M, Legare M, Jenkins C, Rothermel B, et al. Seasonal variation in eastern indigo snake (*Drymarchon couperi*) movement patterns and space use in peninsular Florida at multiple temporal scales. Herpetologica. 2016;72: 214–226.

39. Shamblin BM, Alstad TI, Stevenson DJ, Macey JN, Snow FH, Nairn CJ. Isolation and characterization of microsatellite markers from the threatened eastern indigo snake (*Drymarchon couperi*). Conserv Genet Resour. 2011;3: 303–306. doi:10.1007/s12686-010-9348-5

40. Chapuis MP, Estoup A. Microsatellite null alleles and estimation of population differentiation. Mol Biol Evol. 2007;24: 621–631. doi:10.1093/molbev/msl191

41. Pritchard JK, Stephens M, Donnelly P. Inference of population structure using multilocus genotype data. Genetics. 2000;155: 945–959. doi:10.1111/j.1471-8286.2007.01758.x

42. Earl D, VonHoldt B. STRUCTURE HARVESTER: a website and program for visualizing STRUCTURE output and implementing the Evanno method. Conserv Genet Resour. 2012;4.

43. Evanno G, Regnaut S, Goudet J. Detecting the number of clusters of individuals using the software STRUCTURE: A simulation study. Mol Ecol. 2005;14: 2611–2620. doi:10.1111/j.1365-294X.2005.02553.x

44. Jakobsson M, Rosenburg N. CLUMPP: a cluster matching and permutation program for dealing with label switching and multimodality in analysis of population structure. Bioinformatics. 2007;23: 1801–1806.

45. Janes JK, Miller JM, Dupuis JR, Malenfant RM, Gorrell JC, Cullingham CI, et al. The K = 2 conundrum. Mol Ecol. 2017;26: 3594–3602. doi:10.1111/mec.14187

46. Kopelman N, Mayzel J, Jakobsson M, Rosenberg N, Mayrose I. CLUMPAK: a program for identifying clustering modes and packaging population structure inferences across K. Mol Ecol Resour. 2015;15: 1179–1191.

47. Jost L. GST and its relatives do not measure differentiation. Mol Ecol. 2008;17: 4015– 4026. doi:10.1111/j.1365-294X.2008.03887.x

48. Winter D. MMOD: an R library for the calculation of population differentiation statistics. Mol Ecol Resour. 2012;12.

49. Jombart T, Devillard S, Dufour AB, Pontier D. Revealing cryptic spatial patterns in genetic variability by a new multivariate method. Heredity (Edinb). 2008;101: 92–103. doi:10.1038/hdy.2008.34

50. Stevenson DJ, Hyslop NL. Long-distance interpopulation movement: *Drymarchon couperi* (Eastern Indigo Snake). Herpetol Rev. 2010;41: 91–92.

51. Jombart T. adegenet: a R package for the multivariate analysis of genetic markers. Bioinformatics. 2008;24: 1403–1405.

52. van Strien MJ, Keller D, Holderegger R. A new analytical approach to landscape genetic modelling: Least-cost transect analysis and linear mixed models. Mol Ecol. 2012;21: 4010–4023. doi:10.1111/j.1365-294X.2012.05687.x

53. Bowcock J, Ruiz-Linares A, Tomfohrde J, Minch E, Kidd J, Cavalli-Sforza L. High resolution of human evolutionary trees with polymorphic microsatellites. Nature. 1994;368: 455–457.

54. Nakagawa S, Schielzeth H. A general and simple method for obtaining *R^2^* from generalized linear mixed-effects models. Methods Ecol Evol. 2013;4: 133–142.

55. Johnson P. Extension of Nakagawa & Schielzeth’s *R^2^_GLMM_* to random slopes models. Methods Ecol Evol. 2014;5: 944–946.

56. Peterman W. ResistanceGA: An R package for the optimization of resistance surfaces using genetic algorithms. Methods Ecol Evol. 2018;9: 1638–1647.

57. Ruane S, Bryson RW, Pyron RA, Burbrink FT. Coalescent species delimitation in Milksnakes (Genus *Lampropeltis*) and impacts on phylogenetic comparative analyses. Syst Biol. 2014;63: 231–250. doi:10.1093/sysbio/syt099

58. Meirmans PG. The trouble with isolation by distance. Mol Ecol. 2012;21: 2839–2846. doi:10.1111/j.1365-294X.2012.05578.x

59. Noss RF, Platt WJ, Sorrie BA, Weakley AS, Means DB, Costanza J, et al. How global biodiversity hotspots may go unrecognized: Lessons from the North American Coastal Plain. Divers Distrib. 2015;21: 236–244. doi:10.1111/ddi.12278

60. Burbrink FT, Guiher TJ. Considering gene flow when using coalescent methods to delimit lineages of North American pitvipers of the genus *Agkistrodon*. Zool J Linn Soc. 2014;173: 505–526.

61. Pyron RA, Hsieh FW, Lemmon AR, Lemmon EM, Hendry CR. Integrating phylogenomic and morphological data to assess candidate species-delimitation models in brown and red-bellied snakes (*Storeria*). Zool J Linn Soc. 2016;177: 937–949. doi:10.1111/zoj.12392

62. Mckelvy AD, Burbrink FT. Ecological divergence in the yellow-bellied kingsnake (*Lampropeltis calligaster*) at two North American biodiversity hotspots. Mol Phylogenet Evol. 2017;106: 61–72.

63. Soltis DE, Morris AB, McLachlan JS, Manos PS, Soltis PS. Comparative phylogeography of unglaciated eastern North America. Mol Ecol. 2006;15: 4261–93. doi:10.1111/j.1365-294X.2006.03061.x

64. Noss R. Forgotten grasslands of the South: natural history and conservation. Washington, DC: Island Press; 2013.

65. Strickland JL, Parkinson CL, McCoy JK, Ammerman LK. Phylogeography of *Agkistrodon piscivorus* with Emphasis on the Western Limit of Its Range. Copeia. 2014;2014: 639–649. doi:10.1643/CG-13-123

66. Thomas TM, Granatosky MC, Bourque JR, Krysko KL, Moler PE, Gamble T, et al. Taxonomic assessment of Alligator Snapping Turtles (Chelydridae: *Macrochelys*), with the description of two new species from the southeastern United States. Zootaxa. 2014;3786: 141–165.

67. Echelle AA, Hackler JC, Lack JB, Ballard SR, Roman J, Fox SF, et al. Conservation genetics of the alligator snapping turtle: cytonuclear evidence of range-wide bottleneck effects and unusually pronounced geographic structure. Conserv Genet. 2010;11: 1375– 1387. doi:10.1007/s10592-009-9966-1

68. Folt B, Guyer C. Evaluating recent taxonomic changes for alligator snapping turtles (Testudines: Chelydridae). Zootaxa. 2015;3947: 447–450.

69. Gordon MR, Simandle ET, Tracy CR. A diamond in the rough desert shrublands of the Great Basin in the Western United States: A new cryptic toad species (Amphibia: Bufonidae: *Bufo* (*Anaxyrus*)) discovered in Northern Nevada. Zootaxa. 2017;4290: 123– 139. doi:10.11646/zootaxa.4290.1.7

70. Forrest M, Stiller J, King T, Rouse G. Between hot rocks and dry places: the status of the Dixie Valley Toad. West North Am Nat. 2017;77: 162–175.

71. Madsen T, Shine R. Do snakes shrink? Oikos. 2001;92: 187–188. doi:10.1034/j.1600-0706.2001.920122.x

72. Hyslop NL, Meyers J, Cooper R, Stevenson D. Effects of body size and sex of *Drymarchon couperi* (Eastern Indigo Snake) on habitat use, movements, and home range size in Georgia. J Wildl Manage. 2014;78: 101–111.

73. Breininger DR, Bolt MR, Legare ML, Drese JH, Stolen ED. Factors Influencing Home-Range Sizes of Eastern Indigo Snakes in Central Florida. J Herpetol. 2011;45: 484–490.

74. Bauder JM, Breininger DR, Bolt MR, Legare ML, Jenkins CL, Rothermel BB, et al. The influence of sex and season on conspecific spatial overlap in a large, actively-foraging colubrid snake. PLoS One. 2016;11: 1–19. doi:10.1371/journal.pone.0160033

75. Stiles JA. Evaluating the use of enclosures to reintroduce Eastern Indigo snakes. Auburn University. 2013.

76. Bauder JM, Breininger DR, Bolt MR, Legare ML, Jenkins CL, Rothermel BB, et al. Multi-level, multi-scale habitat selection by a wide-ranging, federally threatened snake. Landsc Ecol. Springer Netherlands; 2018; doi:10.1007/s10980-018-0631-2

77. O’Bryan CJ. Documentation of unusual movement behaviour of the indigo snake *Drymarchon couperi* (Holbrook, 1842) (Squamata: Colubridae), an upland species, in a pastureland matrix of the USA. Herpetol Notes. 2017;10: 317–318.

78. Frankham R, Ballou JD, Eldridge MDB, Lacy RC, Ralls K, Dudash MR, et al. Predicting the Probability of Outbreeding Depression. Conserv Biol. 2011;25: 465–475. doi:10.1111/j.1523-1739.2011.01662.x

79. Thorpe RS, Surget-Groba Y, Johansson H. The relative importance of ecology and geographic isolation for speciation in anoles. Philos Trans R Soc B Biol Sci. 2008;363: 3071–3081. doi:10.1098/rstb.2008.0077

80. Bowen BW, Bass AL, Soares L, Toonen RJ. Conservation implications of complex population structure: Lessons from the loggerhead turtle (*Caretta caretta*). Mol Ecol. 2005;14: 2389–2402. doi:10.1111/j.1365-294X.2005.02598.x

81. de Fraga R, Lima AP, Magnusson WE, Ferrao M, Stow AJ. Contrasting patterns of gene flow for Amazonian snakes that actively forage and those that wait in ambush. J Hered. 2017; 1–11. doi:10.1093/jhered/esx051

82. Leturque H, Rousset F. Intersexual competition as an explanation for sex-ratio and dispersal biases in polygynous species. Evolution (NY). 2004;58: 2398–2408.

83. Keogh J, Webb J, Shine R. Spatial genetic analysis and long-term mark-recapture data demonstrate male-biased dispersal in a snake. Biol Lett. 2007;3: 33–35.

84. Dubey S, Brown GP, Madsen T, Shine R. Male-biased dispersal in a tropical Australian snake (*Stegonotus cucullatus*, Colubridae). Mol Ecol. 2008;17: 3506–3514. doi:10.1111/j.1365-294X.2008.03859.x

85. Calsbeek R. Sex-specific adult dispersal and its selective consequences in the brown anole, *Anolis sagrei*. J Anim Ecol. 2009;78: 617–624. doi:10.1111/j.1365-2656.2009.01527.x

86. Krysko K, Smith D, Smith C. Historic and current geographic distribution and preliminary evidence of population genetic structure in the Eastern Indigo Snake (Drymarchon couperi) in the southeastern United States. Tallahassee, Florida; 2010.

87. Stevenson DJ, Dyer KJ, Willis-Stevenson B a. Survey and Monitoring of the Eastern Indigo Snake in Georgia. Southeast Nat. 2003;2: 393–408. doi:10.1656/1528-7092(2003)002[0393:SAMOTE]2.0.CO;2

88. Hyslop NL, Cooper R, Meyers J. Seasonal shifts in shelter and microhabitat use of the threatened Eastern Indigo Snake (*Drymarchon couperi*) in Georgia. Copeia. 2009;2009: 458–464.

89. Perry G, Garland T. Lizard home ranges revisited: effects of sex, body size, diet, habitat, and phylogeny. Ecology. 2002;83: 1870–1885.

90. Warwick AR, Travis J, Lemmon EM. Geographic variation in the Pine Barrens Treefrog (*Hyla andersonii*): Concordance of genetic, morphometric, and acoustic signal data. Mol Ecol. 2015; n/a-n/a. doi:10.1111/mec.13242

91. Sovic M, Fries A, Gibbs H. Origin of a cryptic lineage in a threatened reptile through isolation and historical hybridization. Heredity (Edinb). 2016;117: 358–366.

92. Macartney JM, Gregory PT, Larsen KW. A tabular survey of data on movements and home ranges of snakes. J Herpetol. 1988;22: 61–73.

